# Label-free deep learning-based species classification of bacteria imaged by phase-contrast microscopy

**DOI:** 10.1101/2023.05.14.540740

**Authors:** Erik Hallström, Vinodh Kandavalli, Petter Ranefall, Johan Elf, Carolina Wählby

## Abstract

Reliable detection and classification of bacteria and other pathogens in the human body, animals, food, and water is crucial for improving and safeguarding public health. For instance, identifying the species and its antibiotic susceptibility is vital for effective bacterial infection treatment. Here we show that phase contrast time-lapse microscopy combined with deep learning is sufficient to discriminate four species of bacteria relevant to human health. The classification is performed on living bacteria and does not require fixation or staining, meaning that the bacterial species can be determined as the bacteria reproduce in a microfluidic device, enabling parallel determination of susceptibility to antibiotics. We assess the performance of convolutional neural networks and vision transformers, where the best model attained a class-average accuracy exceeding 98%. Our successful proof-of-principle results suggest that the methods should be challenged with data covering more species and clinically relevant isolates for future clinical use.

**Author Summary:** Bacterial infections are a leading cause of premature death worldwide, and growing antibiotic resistance is making treatment increasingly challenging. To effectively treat a patient with a bacterial infection, it is essential to quickly detect and identify the bacterial species and determine its susceptibility to different antibiotics. Prompt and effective treatment is crucial for the patient’s survival. A *microfluidic* device functions as a miniature “lab-on-chip” for manipulating and analyzing tiny amounts of fluids, such as blood or urine samples from patients. Microfluidic chips with chambers and channels have been designed for quickly testing bacterial susceptibility to different antibiotics by analyzing bacterial growth. Identifying bacterial species has previously relied on killing the bacteria and applying species-specific fluorescent probes. We introduce deep learning models as a fast and cost-effective method for identifying bacteria species directly from phase-contrast microscopy images of living bacteria simultaneously as growth is analyzed. We envision this method being employed concurrently with antibiotic susceptibility tests in future applications, significantly enhancing bacterial infection treatments.

## 1 Introduction

This study employs deep-learning techniques for species classification of the bacteria *Enterococcus faecalis*, *Escherichia coli*, *Klebsiella pneumoniae*, and *Pseudomonas aeruginosa* cultivated within traps of a microfluidic chip. Combining deep learning methods for data analysis and microfluidics as a data-generating platform has recently spurred significant advances in biotechnology and biomedical research [1]. The widespread success of deep learning across various data-driven fields in recent years has motivated researchers to apply such methods in detecting microbes across various microscopy modalities [2].

The pivotal moment which started the deep learning revolution is commonly accepted to be the development of “AlexNet” by Krizhevsky et al. [3], a deep convolutional neural network (ConvNet) that outperformed competitors in the ImageNet Large Scale Visual Recognition Challenge (ILSVRC2012) in 2012 by a large margin [4]. The success of ConvNets is generally attributed to their ability to automatically learn to extract features through sequential processing of the input data [5].

In the next few years, several techniques and architectural developments further refined the performance of deep convolutional networks [6] [7]. The ResNet (Residual Network) [8] architecture and its variants are currently the primary models utilized in the field, enabling the construction of deeper ConvNets with more layers. ResNet was introduced by He et al. [8], winning the ILSVRC competition in 2015.

More recently, Vision Transformers (ViT) from 2020, a completely novel neural network architecture containing no convolutional filters, has shown to be on par and even surpass convolutional neural networks in various image-processing tasks, including image classification [9]. A transformer is a neural network design initially conceived for sequence-to-sequence modeling in natural language processing tasks, such as language translation or chat robots, first demonstrated in the “Attention is all you need” paper by Vaswani et al. [10].

Various three-dimensional ResNet variants have been used for video clip classification, merging temporal and spatial information across frames [11] [12]. However, recent developments show that transformer-based classifiers considerably surpass these ConvNet methodologies [13].

For this study, we utilized data from Kandavalli et al. [14], where deep learning methods were applied for segmenting and tracking cells growing in a microfluidic chip imaged by phase contrast microscopy. After each completed time-lapse experiment, Kandavalli et al. applied species-specific Fluorescence In Situ Hybridization (FISH) probes and identified bacterial species using images captured by fluorescence microscopy. We leverage the same type of fluorescence microscopy image data as ground truth. However, we use one fluorescent channel per species and do not apply combinatorial FISH [14].

The overall experimental setup is shown in Fig 1. Fig 1A-B: A mixed species sample is loaded into the microfluidic chip. Fig 1C: A phase-contrast time-lapse is captured, recording the growth and reproduction of bacteria in the traps for one hour, consisting of approximately 32 frames. Fig 1C: After fixation and staining, species-specific fluorescent probes attach to each bacteria, and fluorescence microscopy reveals the species. Traps containing only one species are cropped from the phase-contrast timelapse and labeled according to the fluorescent signal. The cropping targets and labels are shown as colored rectangles. Fig 1D: An image or video classification neural network is trained to classify a single frame or time-lapse of growth respectively in a single trap solely using phase-contrast images. Only two out of approximately 200 positions in the microfluidic chip are shown in Fig 1, where half of the positions in an experiment were treated with antibiotics. The reason for including image data of both treated and untreated cells in the pipeline was twofold; firstly, to investigate the robustness of the classification methods, as treated cells can show very different phenotypic characteristics and cell morphologies, and secondly, to probe a possible clinical application where antibiotic susceptibility testing and species classification could be performed simultaneously. Antibiotic susceptibility testing (AST) is performed in a microfluidic device by measuring the differences in the growth rate of treated versus untreated cells to determine how much the cells are affected by antibiotics.

**Fig 1.**
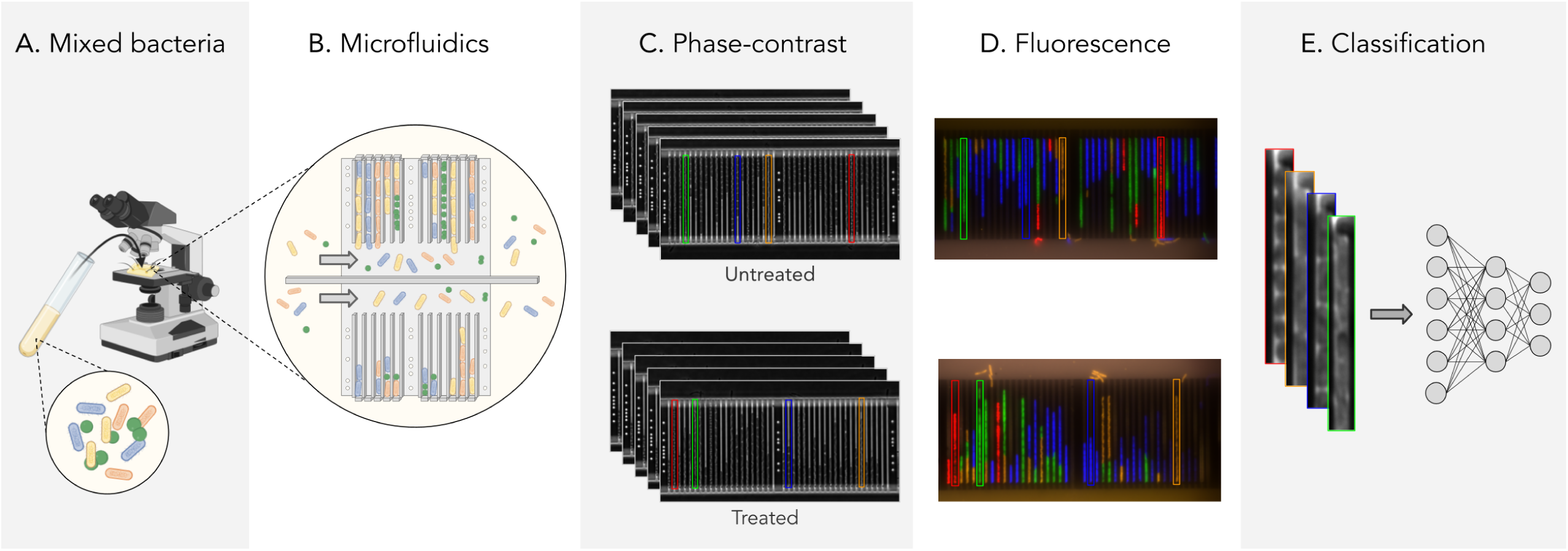
Experiment setup. A, B: An illustration of a mixed bacterial sample loaded on a microfluidic chip. C. Time-lapse phase contrast images of bacterial species growing and reproducing in a medium with (bottom) and without (top) antibiotics. D: The correspondence of the fluorescence image to the final phase-contrast image reveals the species, with *Enterococcus faecalis* (red), *Escherichia coli* (green), *Klebsiella pneumoniae* (blue), and *Pseudomonas aeruginosa* (orange). E: Input to the neural network is a single frame or time-lapse video.

This study explores the potential of using a microfluidic as a viable device for diagnosing the species causing a bacterial infection using deep learning image or video classification methods on phase-contrast image data. Additionally, we aim to carry out species identification in parallel with antibiotics susceptibility testing (AST) on the same microfluidic chip and identify the species as early as possible, eliminating the need for pre-cultivation or staining. Our ultimate goal is integrating species identification with AST to quickly select appropriate patient treatments. To the best of our knowledge, this is the first time ConvNets and Vision Transformers have been used to classify the species of bacteria growing in microfluidic chip traps using only phase-contrast microscopy time-lapses.

## 2 Related work

The authors were unable to find any existing similar studies performing bacteria classification on phase-contrast time lapses. However, several other methods exist for imaging bacteria, such as bright-field, phase-contrast, fluorescence, and electron microscopy at various magnification levels, as exemplified in Zhang et al. [15]. Microscopy images can also be enhanced using different staining methods for better identification and classification of bacteria. Below is a selection of related studies where convolutional neural networks were employed for bacteria classification in micrographs.

Wang et al. [16] developed a system for detecting and classifying live bacteria growing on agar plates. The device captured image data using coherent microscopy, scanning the plate every 30 minutes with an image resolution of 4 µm, generating time-lapses. Two deep neural networks were used, one for the early detection of bacterial growth and the second to classify the species of the bacteria using spatiotemporal data. The classification network could classify three species of bacteria with the accuracies 97.2% for *Escherichia coli*, 84.0% for *Klebsiella aerogenes*, and 98.5% for *Klebsiella pneumoniae* respectively. A custom convolutional neural network design was used for both tasks, using a Pseudo 3D-network [17] coupled with Dense layers [18].

Zieliński et al. [19] performed deep learning classification of 33 bacterial species imaged by bright-field microscopy at 100x magnification. The dataset was released as the DIBaS dataset containing 20 images per species. ConvNet backbones from AlexNet, VGG-M, and VGG-VD pretrained on ImageNet alongside SIFT descriptors were used for feature extraction. Final species classification was performed on the feature representation using either Support Vector Machines or Random forests. Furthermore, an experiment was conducted where several classifiers were trained by increasing the number of species included, measuring the accuracy as a function of the number of classes admitted. The classifiers based on ConvNet feature extraction were shown to have better accuracy than those using SIFT, and the best models acquired a class-average accuracy of around 96%.

Mai et al. [20] further investigated the DIBaS dataset and developed a more efficient classification ConvNet, better tailored for utilization on resource-limited devices. The network used depth-wise separable convolutions, which consist of a depth-wise convolution with one convolutional filter for each input channel, followed by a point-wise 1×1 convolution transforming the input to a desired channel depth. The accuracy was measured using 5-fold cross-validation and revealed a performance almost on par with Zieliński et al. [19] despite using only 3.2 million parameters, significantly less than the heavy backbones pretrained on ImageNet. Rotation, shifting, shearing, scaling, and flip augmentations were demonstrated to be integral to achieving optimal performance. Notably, the absence of these augmentations led to a significant decrease in accuracy.

Smith et al. [21] used convolutional neural networks for automated Gram stain classification. Microscopy image data was captured using a 40x dry objective, and the images were then cropped and annotated manually. In total, 100,213 crops were collected containing Gram-positive cocci in clusters, Gram-positive cocci in chains/pairs, Gram-negative rods or background. The Inception v3 convolutional neural network pretrained on ImageNet was applied to the classification task, and the resulting model attained an average classification accuracy of 94.9% on the held-out test crops, consisting of 20% randomly selected samples from the total crops.

All of the above studies relied on the cultivation of bacteria prior to the classification of bacterial colonies, increasing the time required between isolation of bacteria and final classification. In the presented study, we instead aim to identify bacterial species directly after isolation while growing in a microfluidic chip.

Hay et al. [22] conducted a study in which fluorescently labeled bacteria inside the larval zebrafish gut were imaged with 3D light sheet fluorescence microscopy. Pixel arrays with suspected bacterial content, possibly containing a bacteria cell, were extracted from this image data and independently labeled by six researchers. The 3D pixel arrays had a size of 28×28×8 pixels, with a resolution of 6.22 px/µm, 6.22 px/µm, and 1.0 px/µm, respectively. Ultimately, a ConvNet, a random forest, and a support vector machine (SVM) were trained to classify the pixel arrays automatically, whereas the last two methods utilized texture-based feature extraction. The ConvNet reached near-human accuracy and was shown to outperform the other methods with both accuracy and inference speed. Furthermore, transfer learning and augmentation were demonstrated to be highly beneficial for classification accuracy. Compared to the presented study, this study relied on fluorescently labeled cells, making it infeasible for clinical samples.

Panigrahi et al. [23] used a shape index map calculated from the Hessian of the image as a preprocessing step before feeding the single-channel phase-contrast image data to a U-net for semantic classification of *Myxococcus xanthus* and *Escherichia coli*. Genetically modified bacteria expressing fluorescent markers (GFP and mCherry for *Myxococcus xanthus* and *Escherichia coli*, respectively) were used to generate training data. The image data had a resolution of 16.67 px/µm. The authors reported Jaccard Index test scores of 0.95 ± 0.036 (n = 200 cells) and 0.89 ± 0.047 (n = 545 cells) for the two bacterial species, respectively. This approach is limited to classifying two bacterial species with pronounced differences in size and shape under unconstrained growth conditions. Additionally, this technique is incompatible with antibiotic susceptibility testing. In the work presented in this paper, we classify four bacterial species of similar size, leveraging further information such as texture and cell interaction during growth inside a microfluidic chip trap. Furthermore, we employ video classification methods to extract temporal data for more accurate species identification.

To conclude, the advantage of using phase-contrast micrographs from a microfluidic chip is that no pre-cultivation or staining is required, and cropping out traps can be automated, resulting in faster inference speed. In a clinical setting, it can be performed “on the fly” by continuously acquiring more time-lapse video data as time progresses, increasing the prediction’s confidence in real time, and also enabling parallel antibiotics susceptibility testing. Furthermore, the novel Vision Transformer and the video classification network “Video ResNet” is particularly robust during inference when using subsampled image data, making it compatible with lower magnification microscopy and suitable for future clinical implementation.

## 3 Results

In our study, we assessed the classification accuracy of three types of deep-learning models: Vision Transformers (ViT), ResNets, and R(2+1)D “Video ResNets” [11]. The ViT and ResNets were investigated using varying model sizes and sub-sampled input image data. The Video ResNets were evaluated by feeding the network single or multiple time-lapse frames. Additionally, experiments were conducted using spatially subsampled time-lapses processed by the Video ResNet.

The models were trained and evaluated on 3,396 cropped-out time-lapse videos of single traps with bacteria growing in a microfluidic chip. Each video contained 32 image frames collected at a frame rate of two minutes. The image time-lapse data were collected from several experiments where all four bacterial species were mixed and cultured in the same microfluidic device. In addition to time-lapses of untreated bacteria, the dataset included bacteria treated with different antibiotics (see Materials and Methods). These mixed species datasets were selected to avoid potential classification bias arising from chip-to-chip variations. Some traps contained a mix of bacteria from different species, visible from the fluorescence signals captured after the final frame. These traps were excluded from the dataset as the probability of mixed species in the same trap is very low in an actual clinical setting. However, this ground truth selection was occasionally unreliable (see Discussion).

The data were partitioned into train/test by trap-basis so that image data from 85% of the traps were used to train the deep learning models, and 15% of the traps were used for testing. Hence all models were tested on unseen images and time-lapses, and the models could not train and test on images originating from the same trap. We intentionally trained standard models with default settings (augmentation, epochs, batch size, and learning rates) until convergence without adjusting any hyperparameters. Consequently, we did not utilize a validation set.

Each network was retrained five times to obtain more accurate statistics, using a predetermined random seed for each retraining. This approach ensured a reproducible train/test split, augmentation sequence, and weight initialization.

The accuracy was first calculated separately for each of the four species as the number of correctly classified instances of a particular species, true positives, *TP_i_* over the total number of instances of that species *N_i_* in the dataset, 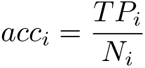, referred to as class-specific or “species-specific” accuracy. The class-specific accuracies were then averaged to obtain the class-average accuracy describing the overall accuracy of a model, 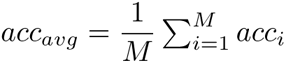, where *M* = 4 is the number of species in the dataset. In the context of multi-class problems, this metric is alternatively labeled as macro-averaged accuracy.

### 3.1 Single-frame classification

We first evaluated our networks’ ability to identify bacterial species solely using the first frame in the time-lapse - this would represent the very first information available in a clinical setting. For this single-frame classification experiment, ResNets and ViTs were trained on all time-lapse frames in the training set but evaluated only on the first frame of the time-lapses in the test set. Networks of different sizes were compared, either trained from scratch or pretrained on ImageNet. The base transformers ViT-B/16 and ViT-B/8 are the standard transformer models from [9] with patch sizes 16 and 8 respectively, the others are downscaled versions of ViT-B/8, with the settings ViT-[*d_embed_*]-[*h*]-[*n_depth_*]. As seen in the results in Fig 2 and Fig 3, the pretrained ViT outperformed the best-achieving ResNet despite the ResNet’s inherent inductive bias for image classification. Pretraining was shown to be more critical for ViT than for ResNets. Deeper networks yielded robust gains, however, the accuracy plateaued when using a ResNet with more than 26 layers. Smaller models, in particular, had a higher tendency for classification errors on *Pseudomonas aeruginosa*.

**Fig 2.**
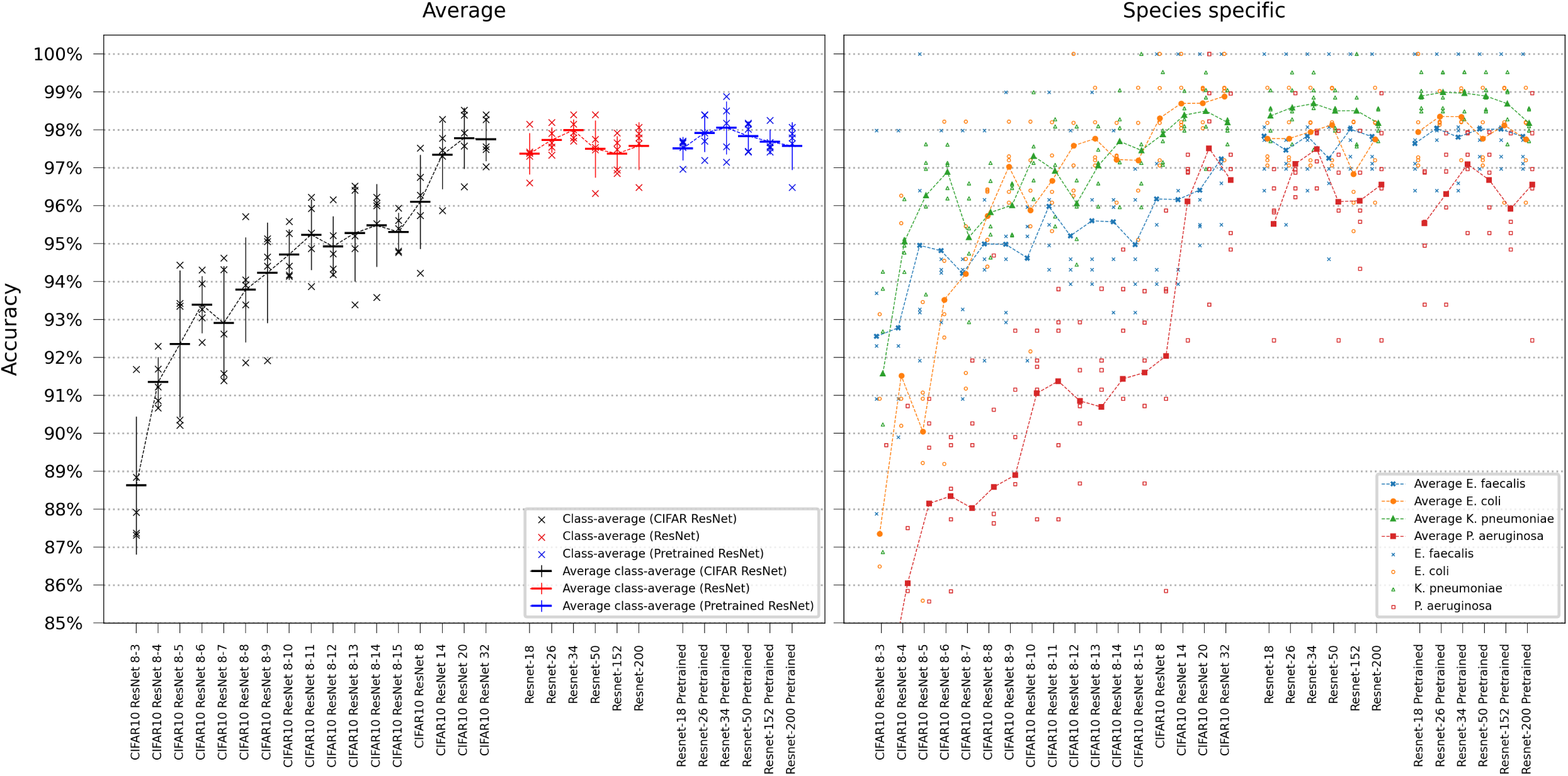
ResNet single-frame classification. Model comparison performing single-frame classification of the first frame in the time-lapse using different ResNet variants. The models are categorized based on the ResNet family. Error bars represent the standard deviation in class-average accuracy from the five retrainings. Scatter plots depict the class-specific accuracy of all individual classifiers and the average class-specific accuracy over the five retrainings for each model. To reduce overplotting, a minor jitter was introduced along the categorical axis of the species-specific scatter plot. Lines are included not for interpolation or statistical inference purposes but to visually guide readers in tracking mean values on the ordinal scale.

**Fig 3.**
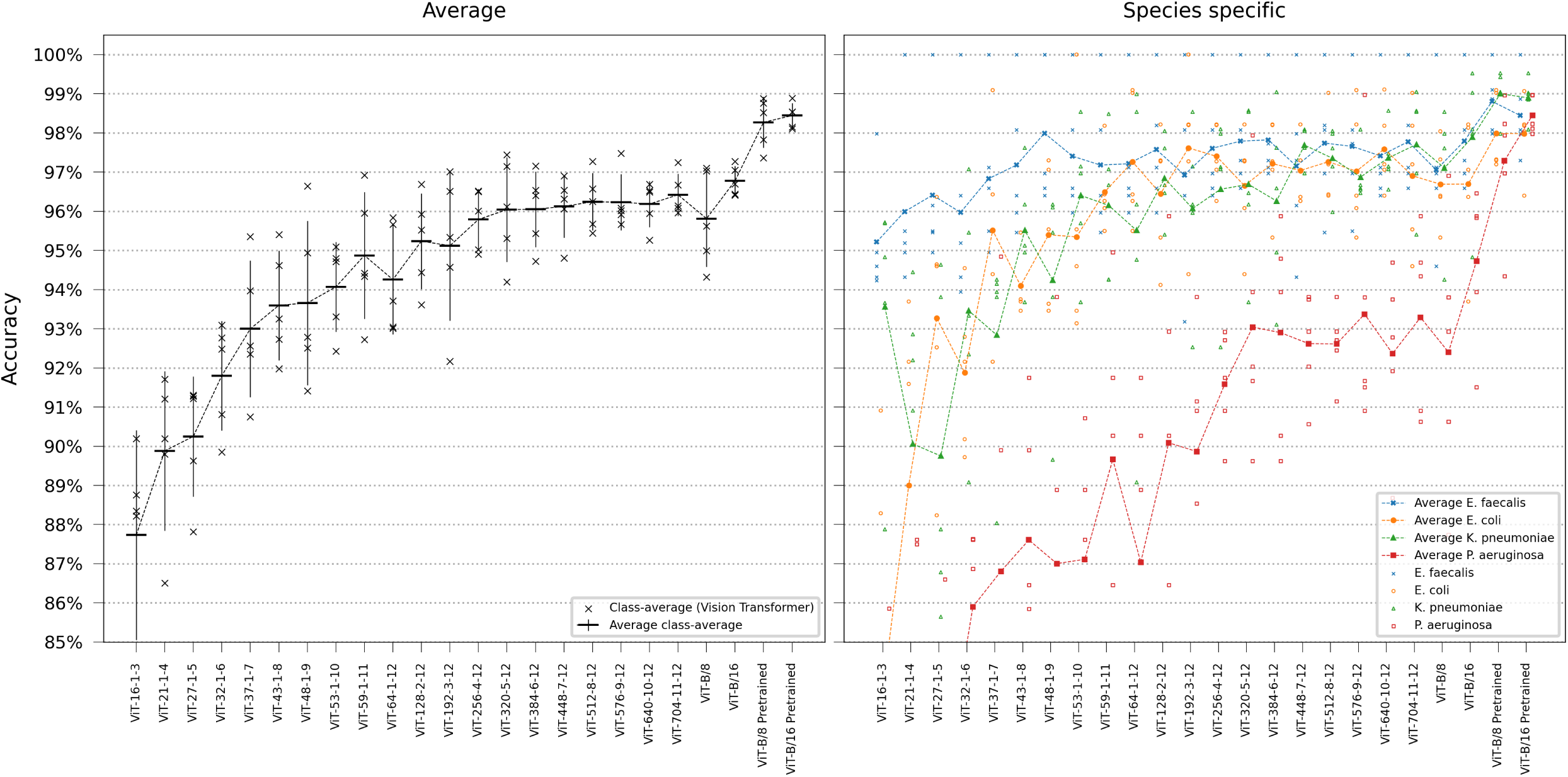
ViT single-frame classification. Model comparison performing single-frame classification of the first frame in the time-lapse using different ViT variants. Error bars represent the standard deviation in class-average accuracy from the five retrainings. Scatter plots depict the class-specific accuracy of all individual classifiers and the average class-specific accuracy over the five retrainings for each model. To reduce overplotting, a minor jitter was introduced along the categorical axis of the species-specific scatter plot. Lines are included not for interpolation or statistical inference purposes but to visually guide readers in tracking mean values on the ordinal scale.

#### 3.1.1 Training and testing on a balanced non-treated dataset

In the single-frame experiments, most classification errors occurred with *Pseudomonas aeruginosa*. This trend was especially noticeable when employing smaller ResNets, non-pretrained downsized Vision Transformers, or during the subsampling experiments. We hypothesized that the challenge arose due to the overrepresentation of treated *Pseudomonas aeruginosa*, with only 250 out of 662 untreated samples available. We suspected these treated traps resulted in altered image conditions, thereby complicating the classification. To test this hypothesis, we conducted additional experiments using the downscaled versions of ResNet-8 and ViT Base patch 8, selecting only 250 traps of each species, reserving 15% of the time-lapses for testing as done previously, and evaluating on the first frame. The results showed similar scaling characteristics detailed further in S1 Appendix.

### 3.1.2 Testing on later frames

Since the assigned class label is determined immediately after the final frame in the time-lapse sequence, subsequent frames may more closely correspond to the actual label. Furthermore, the effects of antibiotic treatment will only be apparent in later stages, leading to alterations in cell appearance (see Discussion). We conducted additional experiments to investigate this, assessing single-frame accuracy testing on later frames in the time-lapse using ResNet-26 and ViT-16. The results are shown in Fig 4. The vision transformer consistently outperforms ResNet and is more accurate with lower variance across the test partitions. There were no evident accuracy changes observed when testing on later frames.

**Fig 4.**
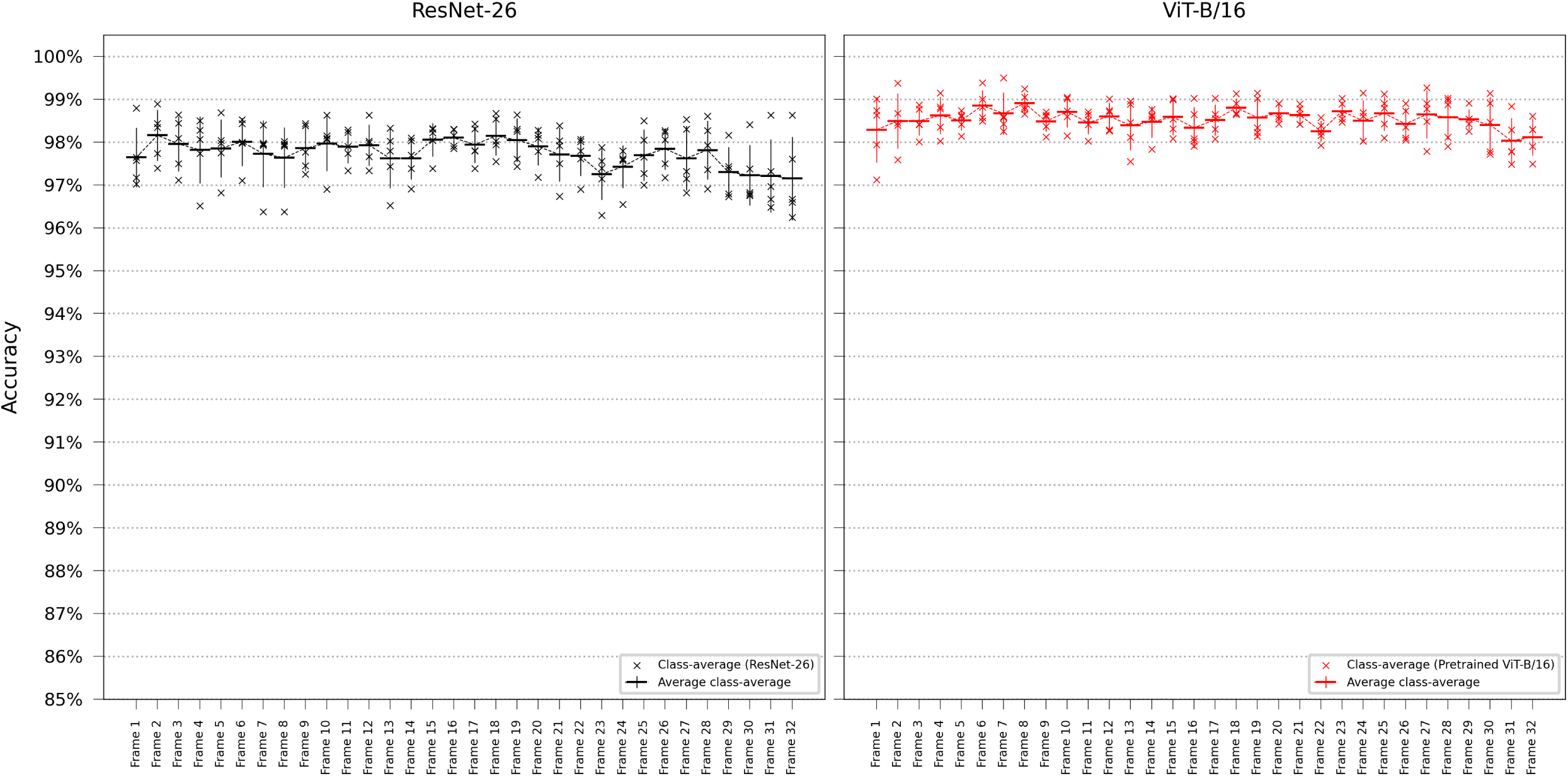
Single-frame classification testing on later frames. Performing single-frame classification, testing on subsequently later frames in the time-lapse using ResNet-26 and ViT-B/16. Error bars represent the standard deviation in class-average accuracy from the five retrainings. Scatter plots depict the class-specific accuracy of all individual classifiers and the average class-specific accuracy over the five retrainings for each model. To reduce overplotting, a minor jitter was introduced along the categorical axis of the species-specific scatter plot. Lines are included not for interpolation or statistical inference purposes but to visually guide readers in tracking mean values on the ordinal scale.

**Fig 5.**
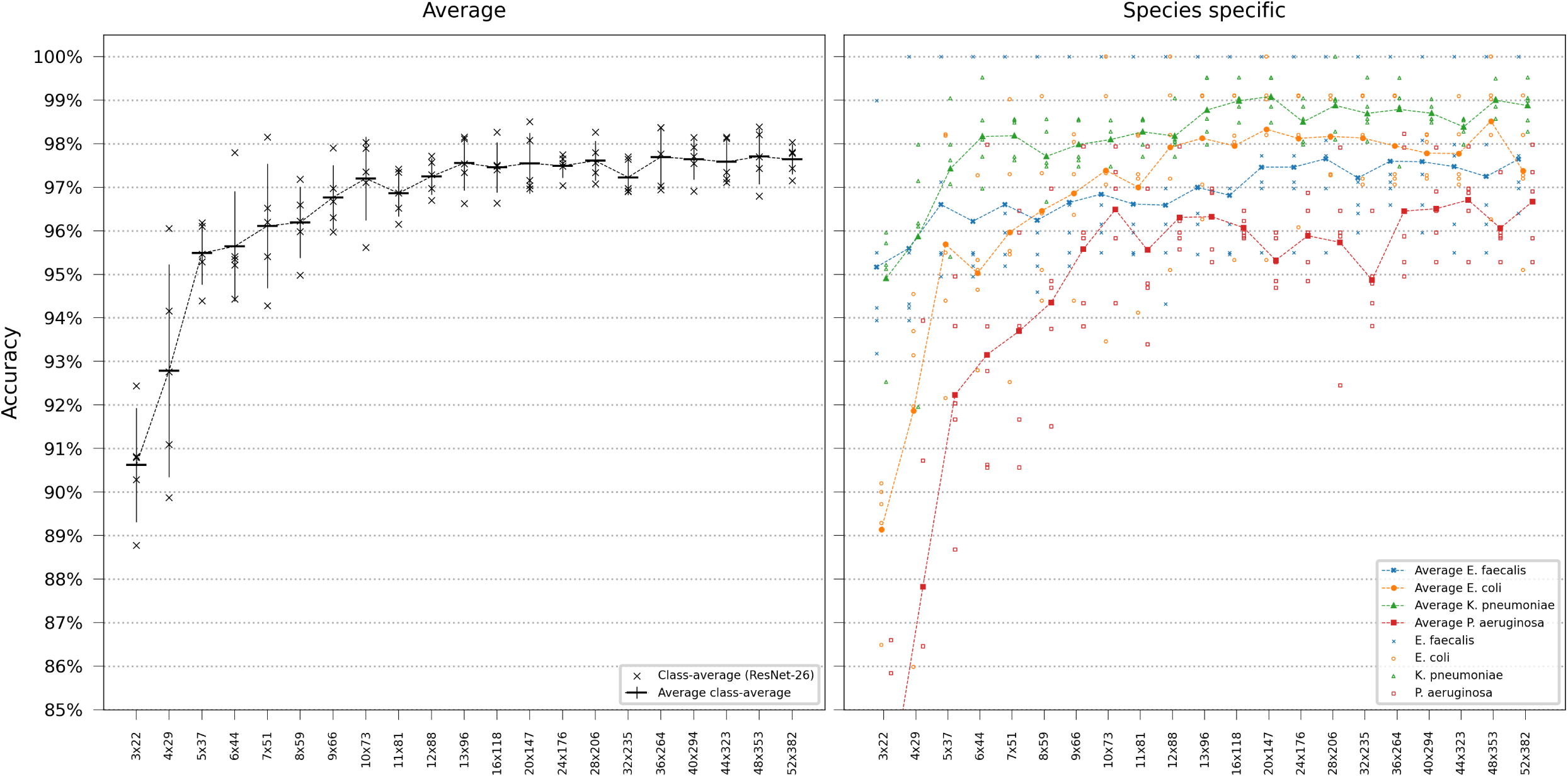
ResNet performance at reduced image resolution. Error bars represent the standard deviation in class-average accuracy from the five retrainings. Scatter plots depict the class-specific accuracy of all individual classifiers and the average class-specific accuracy over the five retrainings for each model. To reduce overplotting, a minor jitter was introduced along the categorical axis of the species-specific scatter plot. Lines are included not for interpolation or statistical inference purposes but to visually guide readers in tracking mean values on the ordinal scale.

## 3.2 Decreasing resolution

Subsequently, we conducted experiments to simulate using lower magnification microscopy to assess the model’s viability in potential clinical devices for two primary reasons: These devices are usually equipped with lower-resolution microscopy and have constrained computational capacities. For these evaluations, we chose the pretrained models ResNet-26 and ViT-B/8, as ViT-B/8 could handle smaller image sizes than ViT-B/16, and deeper ResNets did not notably increase accuracy. We initiated our training using models with the original image dimensions of 52×382 pixels and decreased the resolution step-wise, ending up in training models with an image size of 3×22 pixels. Given that the ViT has a patch size of 8×8 pixels, we were unable to reduce the resolution below a width of 8 pixels for the transformer. Consequently, the evaluation was limited, so the smallest image dimensions were 8×59 pixels. Sub-sampled images from all frames in the time-lapse of the training dataset were used during training, and testing was performed using a sub-sampled version of the first frame in the hold-out test dataset. As illustrated in Fig 8 and 6, ViT-B/8 consistently performed better than ResNet-26 when using sub-sampled images.

**Fig 6.**
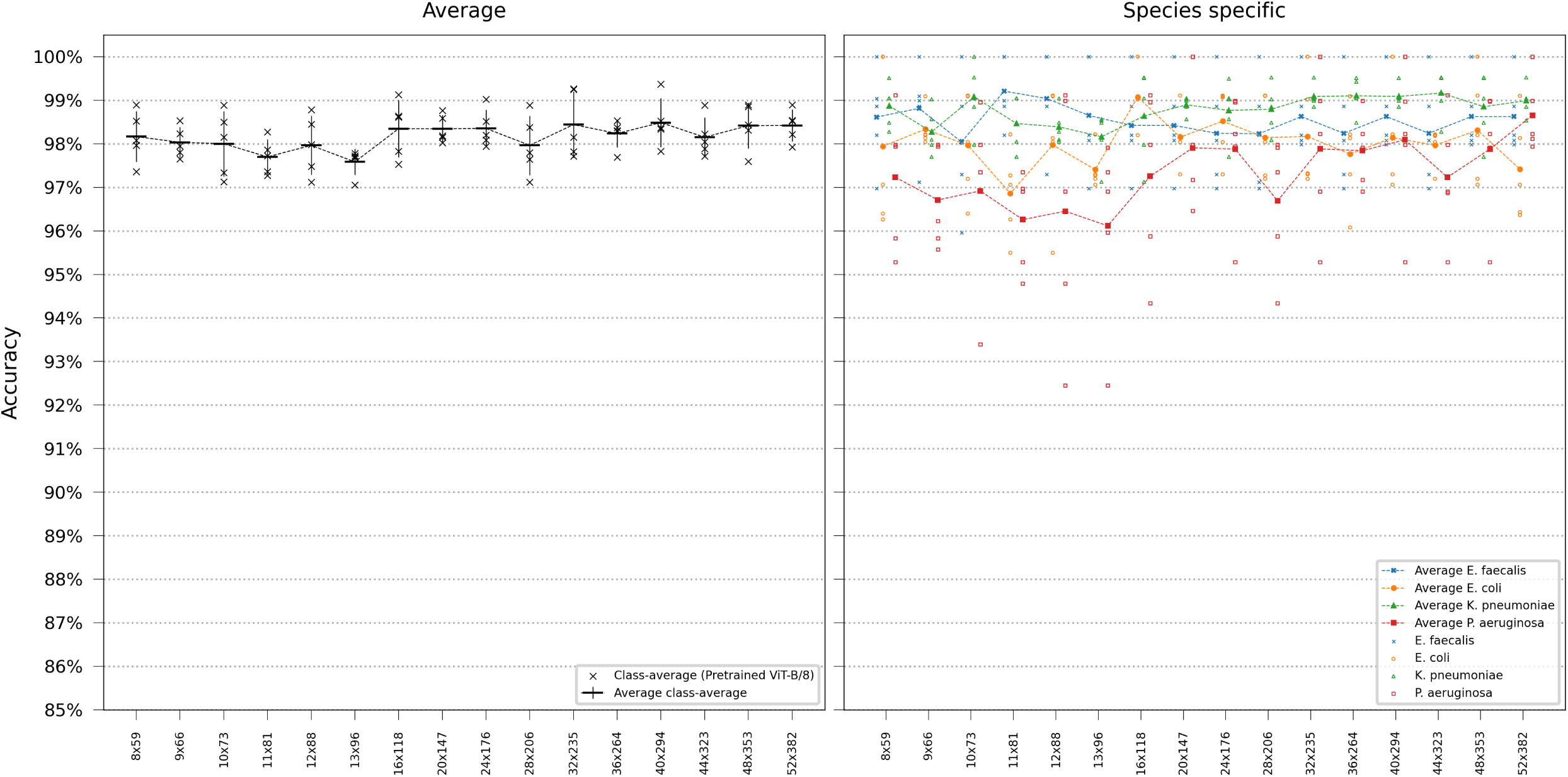
ViT performance at reduced image resolution. Error bars represent the standard deviation in class-average accuracy from the five retrainings. Scatter plots depict the class-specific accuracy of all individual classifiers and the average class-specific accuracy over the five retrainings for each model. To reduce overplotting, a minor jitter was introduced along the categorical axis of the species-specific scatter plot. Lines are included not for interpolation or statistical inference purposes but to visually guide readers in tracking mean values on the ordinal scale.

## 3.3 Video classification

After establishing network performance on single frames, we investigated whether accuracy could be improved using multiple frames and if the network could leverage temporal information from the bacteria reproduction. The R(2+1)D Video ResNet model was trained using an incrementally increasing number of frames as input, with the frames either shuffled or ordered. Testing was performed starting from the first frame in the test time-lapses. As illustrated in Fig 7, accuracy improves as the classifier can access more frames. Furthermore, preserving the original sequence of time-lapse frames results in slightly enhanced performance compared to shuffled frames, particularly when using a larger number of frames.

**Fig 7.**
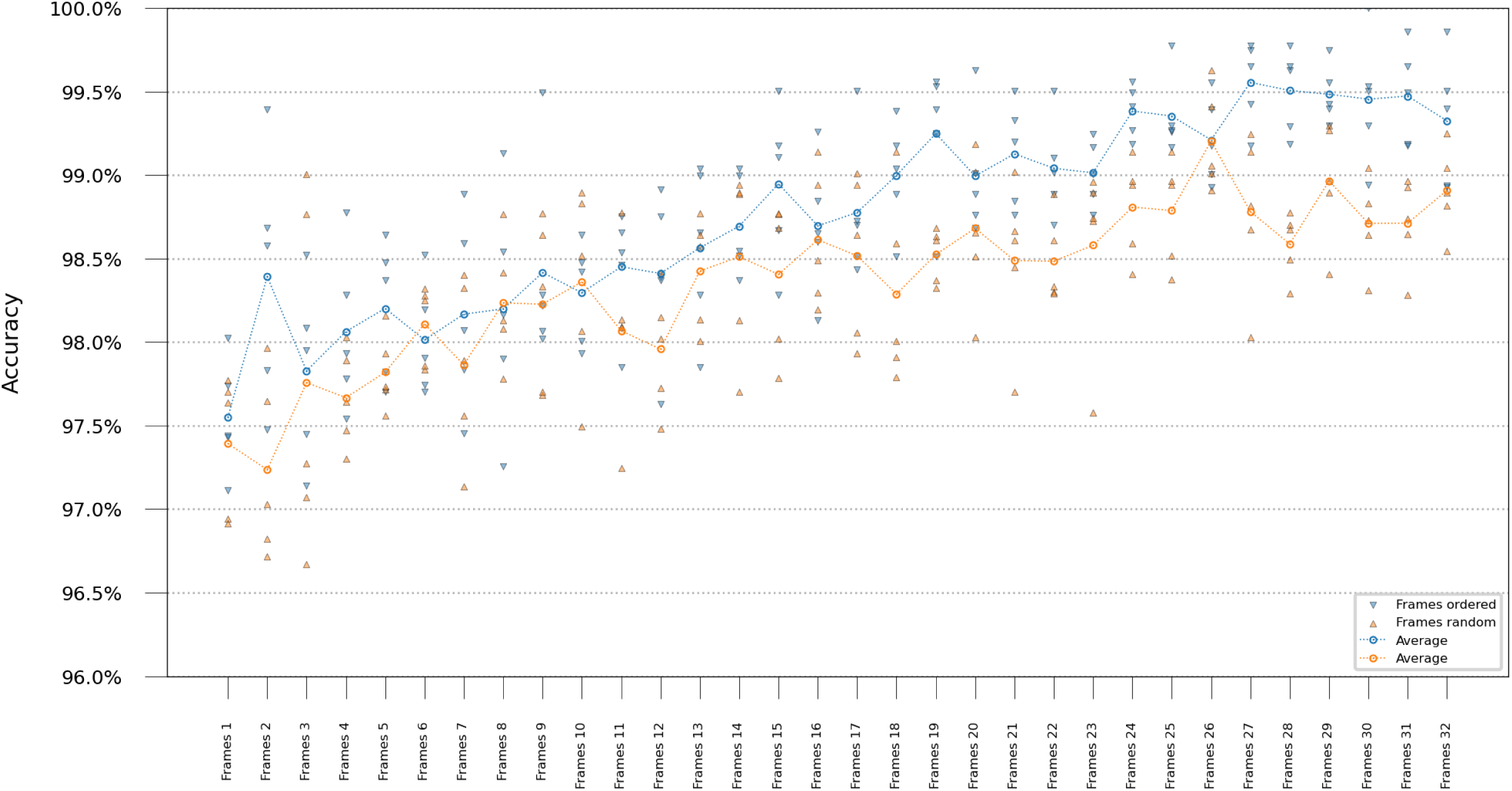
Video ResNet performance at varying number of frames. Results from training video classification “R(2+1)D”-networks using various numbers of frames. Ordered and randomly shuffled time-lapses were compared for a fixed number of frames. The scatter plot shows class-average accuracy for each classifier, the line graph shows average class-average accuracy over all classifiers retrained with their respective train/test split. Lines are not for interpolation or statistical inference but are added as a visual guide to track the mean values across the ordinal scale.

**Fig 8.**
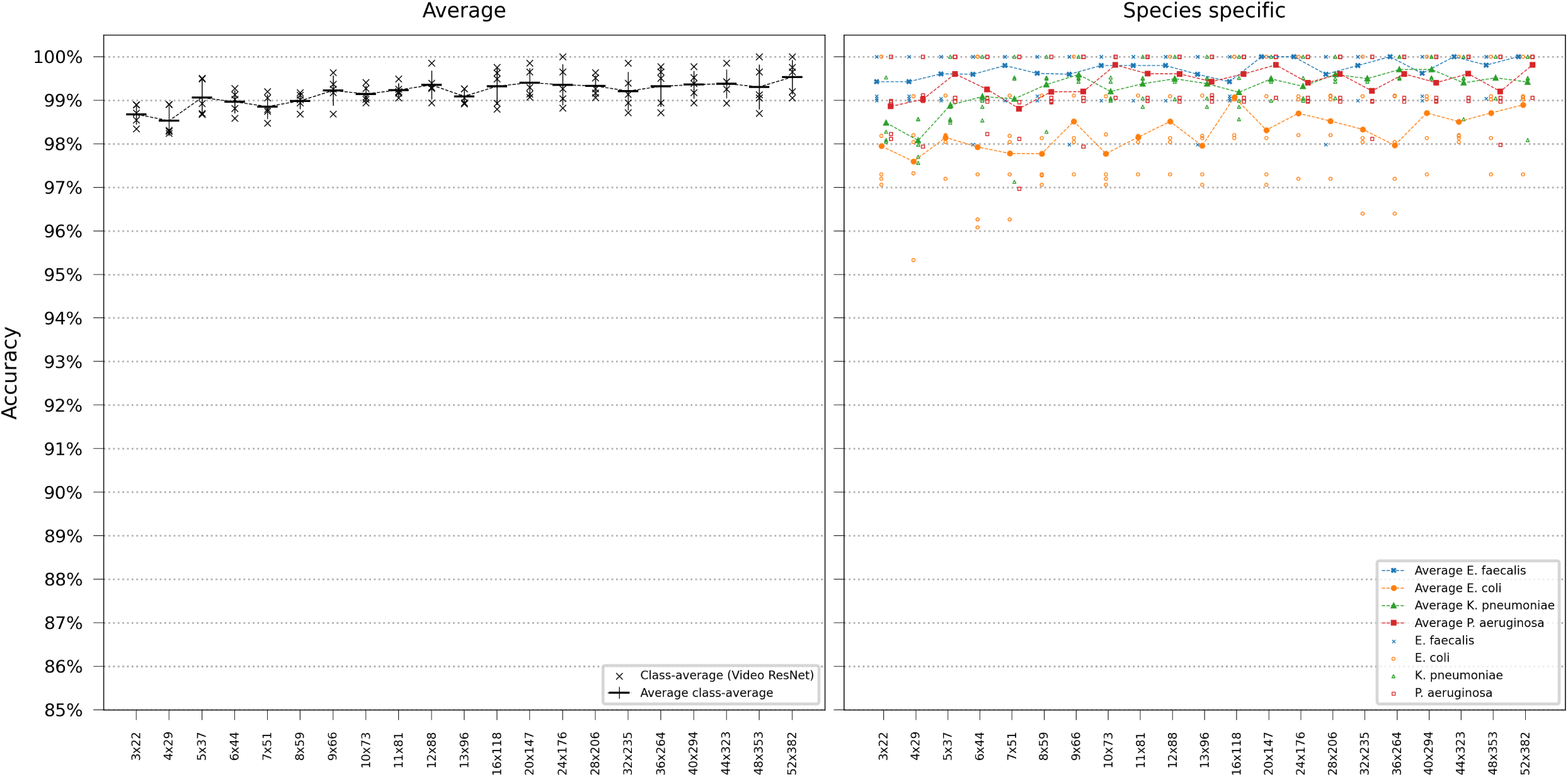
Video ResNet performance at reduced image resolution. Error bars represent the standard deviation in class-average accuracy from the five retrainings. Scatter plots depict the class-specific accuracy of all individual classifiers and the average class-specific accuracy over the five retrainings for each model. To reduce overplotting, a minor jitter was introduced along the categorical axis of the species-specific scatter plot. Lines are included not for interpolation or statistical inference purposes but to visually guide readers in tracking mean values on the ordinal scale.

### 3.3.1 Decreasing resolution

In our final evaluation, we performed video classification employing R(2+1)D Video ResNet, incorporating all frames in the time-lapse, training, and testing on spatially subsampled image data. Notably, even at considerably reduced resolutions, the model retained high accuracy. However, we observed a consistent, albeit slight, decline in accuracy (approximately 1%) as the model was trained and tested on progressively lower resolutions.

## 3.4 Model evaluation

All model results are summarized in Table 1 and 2. The models ResNet-26, ViT-B/8, and 32-frame R(2+1)D Video ResNet from the scaling experiments were selected and trained from scratch using an identical random seed and thus train/test split. Confusion matrices in Fig 9 show classification errors for these models. In order to evaluate model limitations and analyze image data in instances where the model makes an incorrect assessment, the misclassified test samples from the models in Fig 9 are outlined in S1 Fig - S12 Fig.

**Fig 9.**
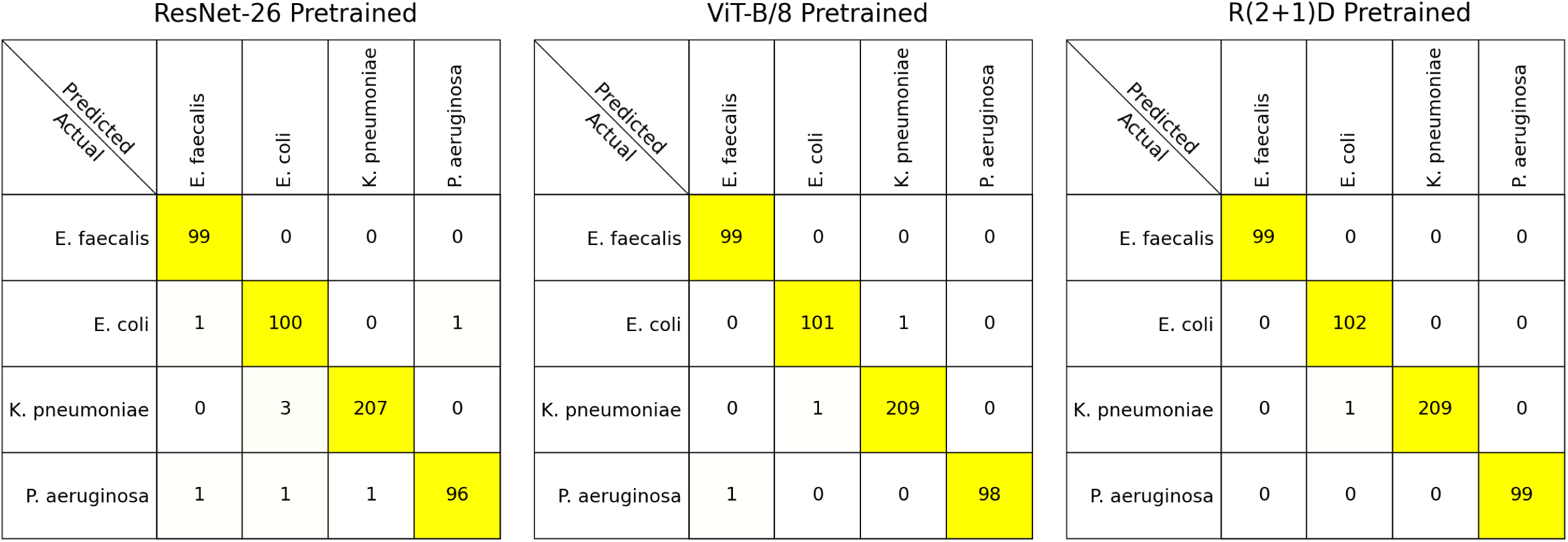
Confusion matrices. Confusion matrices of the models trained once from scratch using the same random seed. All models were trained and tested at full resolution. The Video ResNet “R(2+1)D” used all 32 frames during both the training and testing phases. The single-frame classifiers ResNet-26 and ViT-B/8 trained on all frames in the time-lapse but could only access the first frame during test time.

**Table 1.** Model results: single-frame.

| ResNet |  |  |  | ViT |  |  |  |
| --- | --- | --- | --- | --- | --- | --- | --- |
| Model | Parameters | FLOPS | Accuracy | Model | Parameters | FLOPS | Accuracy |
| CIFAR10 ResNet 8-3 | 2887 | 17.32 MFlops | 88.63 ± 1.82% | ViT-16-1-3 | 15.24 k | 19.21 MFlops | 87.73 ± 2.68% |
| CIFAR10 ResNet 8-4 | 5040 | 29.84 MFlops | 91.35 ± 0.66% | ViT-21-1-4 | 29.34 k | 36.03 MFlops | 89.88 ± 2.04% |
| CIFAR10 ResNet 8-5 | 7789 | 45.71 MFlops | 92.35 ± 1.94% | ViT-27-1-5 | 54.60 k | 62.80 MFlops | 90.25 ± 1.54% |
| CIFAR10 ResNet 8-6 | 11.13 k | 64.96 MFlops | 93.39 ± 0.75% | ViT-32-1-6 | 87.01 k | 95.19 MFlops | 91.80 ± 1.40% |
| CIFAR10 ResNet 8-7 | 15.07 k | 87.58 MFlops | 92.90 ± 1.51% | ViT-37-1-7 | 130.84 k | 136.42 MFlops | 93.00 ± 1.74% |
| CIFAR10 ResNet 8-8 | 19.61 k | 113.56 MFlops | 93.78 ± 1.38% | ViT-43-1-8 | 196.47 k | 194.10 MFlops | 93.60 ± 1.40% |
| CIFAR10 ResNet 8-9 | 24.75 k | 142.91 MFlops | 94.23 ± 1.33% | ViT-48-1-9 | 270.63 k | 257.27 MFlops | 93.66 ± 2.10% |
| CIFAR10 ResNet 8-10 | 30.47 k | 175.63 MFlops | 94.72 ± 0.67% | ViT-53-1-10 | 361.83 k | 332.27 MFlops | 94.07 ± 1.15% |
| CIFAR10 ResNet 8-11 | 36.80 k | 211.71 MFlops | 95.23 ± 0.93% | ViT-59-1-11 | 487.82 k | 431.43 MFlops | 94.87 ± 1.62% |
| CIFAR10 ResNet 8-12 | 43.72 k | 251.17 MFlops | 94.92 ± 0.80% | ViT-64-1-12 | 621.38 k | 534.74 MFlops | 94.25 ± 1.40% |
| CIFAR10 ResNet 8-13 | 51.24 k | 293.99 MFlops | 95.28 ± 1.28% | ViT-128-2-12 | 2.42 M | 1.69 GFlops | 95.24 ± 1.22% |
| CIFAR10 ResNet 8-14 | 59.35 k | 340.18 MFlops | 95.48 ± 1.09% | ViT-192-3-12 | 5.40 M | 3.48 GFlops | 95.12 ± 1.91% |
| CIFAR10 ResNet 8-15 | 68.06 k | 389.74 MFlops | 95.30 ± 0.51% | ViT-256-4-12 | 9.56 M | 5.89 GFlops | 95.79 ± 0.79% |
| CIFAR10 ResNet 8 | 77.36 k | 442.66 MFlops | 96.10 ± 1.24% | ViT-320-5-12 | 14.90 M | 8.93 GFlops | 96.04 ± 1.34% |
| CIFAR10 ResNet 14 | 174.58 k | 952.83 MFlops | 97.34 ± 0.90% | ViT-384-6-12 | 21.42 M | 12.59 GFlops | 96.05 ± 0.97% |
| CIFAR10 ResNet 20 | 271.80 k | 1.46 GFlops | 97.78 ± 0.81% | ViT-448-7-12 | 29.12 M | 16.87 GFlops | 96.12 ± 0.80% |
| CIFAR10 ResNet 32 | 466.23 k | 2.48 GFlops | 97.75 ± 0.57% | ViT-512-8-12 | 38.00 M | 21.78 GFlops | 96.24 ± 0.73% |
| Resnet-18 | 11.17 M | 1.43 GFlops | 97.37 ± 0.55% | ViT-576-9-12 | 48.06 M | 27.32 GFlops | 96.23 ± 0.72% |
| Resnet-26 | 13.95 M | 1.88 GFlops | 97.73 ± 0.33% | ViT-640-10-12 | 59.30 M | 33.48 GFlops | 96.18 ± 0.59% |
| Resnet-34 | 21.28 M | 3.00 GFlops | 97.99 ± 0.28% | ViT-704-11-12 | 71.72 M | 40.27 GFlops | 96.42 ± 0.54% |
| Resnet-50 | 23.51 M | 3.37 GFlops | 97.50 ± 0.75% | ViT-B/8 | 85.31 M | 47.68 GFlops | 95.81 ± 1.23% |
| Resnet-152 | 58.15 M | 9.95 GFlops | 97.37 ± 0.46% | ViT-B/16 | 85.31 M | 11.59 GFlops | 96.77 ± 0.38% |
| Resnet-200 | 62.63 M | 12.71 GFlops | 97.58 ± 0.63% | ViT-B/8 Pretrained | 85.31 M | 47.68 GFlops | 98.27 ± 0.65% |
| Resnet-18 Pretrained | 11.17 M | 1.43 GFlops | 97.50 ± 0.31% | **ViT-B/16 Pretrained** | **85.31 M** | **11.59 GFlops** | **98.44 ± 0.32%** |
| Resnet-26 Pretrained | 13.95 M | 1.88 GFlops | 97.92 ± 0.51% | ResNet-26 Pretrained - Decrease resolution | | | |
| **Resnet-34 Pretrained** | **21.28 M** | **3.00 GFlops** | **98.05 ± 0.70%** | Resolution | Parameters | FLOPS | Accuracy |
| Resnet-50 Pretrained | 23.51 M | 3.37 GFlops | 97.84 ± 0.38% | 3x22 | 13.95 M | 38.53 MFlops | 90.62 ± 1.31% |
| Resnet-152 Pretrained | 58.15 M | 9.95 GFlops | 97.68 ± 0.33% | 4x29 | 13.95 M | 40.81 MFlops | 92.79 ± 2.45% |
| Resnet-200 Pretrained | 62.63 M | 12.71 GFlops | 97.58 ± 0.63% | 5x37 | 13.95 M | 72.41 MFlops | 95.49 ± 0.72% |
| ViT-B/8 Pretrained - Decrease resolution | | | | 6x44 | 13.95 M | 75.43 MFlops | 95.64 ± 1.27% |
| Resolution | Parameters | FLOPS | Accuracy | 7x51 | 13.95 M | 76.36 MFlops | 96.11 ± 1.43% |
| 8x59 | 85.12 M | 1.19 GFlops | 98.16 ± 0.58% | 8x59 | 13.95 M | 85.43 MFlops | 96.19 ± 0.82% |
| 9x66 | 85.12 M | 1.36 GFlops | 98.03 ± 0.35% | 9x66 | 13.95 M | 106.52 MFlops | 96.77 ± 0.74% |
| 10x73 | 85.12 M | 1.53 GFlops | 98.00 ± 0.75% | 10x73 | 13.95 M | 136.72 MFlops | 97.20 ± 0.96% |
| 11x81 | 85.12 M | 1.71 GFlops | 97.70 ± 0.40% | 11x81 | 13.95 M | 142.23 MFlops | 96.86 ± 0.54% |
| 12x88 | 85.12 M | 1.88 GFlops | 97.96 ± 0.68% | 12x88 | 13.95 M | 153.61 MFlops | 97.25 ± 0.42% |
| 13x96 | 85.12 M | 2.05 GFlops | 97.58 ± 0.30% | 13x96 | 13.95 M | 162.94 MFlops | 97.55 ± 0.63% |
| 16x118 | 85.13 M | 4.62 GFlops | 98.35 ± 0.65% | 16x118 | 13.95 M | 205.98 MFlops | 97.46 ± 0.58% |
| 20x147 | 85.14 M | 5.66 GFlops | 98.34 ± 0.32% | 20x147 | 13.95 M | 351.07 MFlops | 97.55 ± 0.70% |
| 24x176 | 85.16 M | 10.52 GFlops | 98.35 ± 0.43% | 24x176 | 13.95 M | 442.43 MFlops | 97.49 ± 0.28% |
| 28x206 | 85.16 M | 12.09 GFlops | 97.96 ± 0.68% | 28x206 | 13.95 M | 539.01 MFlops | 97.61 ± 0.46% |
| 32x235 | 85.19 M | 18.99 GFlops | 98.44 ± 0.76% | 32x235 | 13.95 M | 647.23 MFlops | 97.23 ± 0.41% |
| 36x264 | 85.20 M | 21.13 GFlops | 98.24 ± 0.32% | 36x264 | 13.95 M | 1.05 GFlops | 97.70 ± 0.70% |
| **40x294** | **85.24 M** | **29.26 GFlops** | **98.49 ± 0.56%** | 40x294 | 13.95 M | 1.19 GFlops | 97.64 ± 0.47% |
| 44x323 | 85.25 M | 32.92 GFlops | 98.15 ± 0.45% | 44x323 | 13.95 M | 1.41 GFlops | 97.58 ± 0.51% |
| 48x353 | 85.29 M | 43.15 GFlops | 98.42 ± 0.52% | **48x353** | **13.95 M** | **1.57 GFlops** | **97.70 ± 0.64%** |
| 52x382 | 85.31 M | 47.68 GFlops | 98.42 ± 0.36% | 52x382 | 13.95 M | 1.88 GFlops | 97.64 ± 0.35% |

Average test accuracies and their corresponding standard deviations were computed across five retraining iterations, each utilizing a predetermined random seed.

## 4 Discussion

The findings indicate the feasibility of training deep learning models to accurately classify the bacteria species *E. faecalis*, *E. coli*, *K. pneumoniae*, and *P. aeruginosa* growing in traps in microfluidic chips based solely on phase-contrast image data. Classification accuracy can be maintained at lower image resolution and, thus, lower computational cost, which enables the models to be implemented in potential diagnostics tools utilizing lower-resolution microscopy and equipped with limited computing resources. In the single-frame study, ViTs marginally outperformed ResNets, especially at lower resolutions. The Video ResNet” R(2+1)D” using 27 frames attained the best overall accuracy, surpassing 95.5%. Moreover, when the video classification network was applied to spatially downsampled video data, the model exhibited exceptional performance, successfully classifying even the most low-resolution time-lapses with little accuracy degradation.

The models were pretrained using the ImageNet dataset [4], which predominantly contains natural images, contrasting significantly with microscopy images of bacteria. While pretraining is deemed crucial in conventional settings, particularly for transformers, it was not immediately evident whether pretraining would be beneficial for processing the bacteria image data in this study. However, the results showed that this was critical for transformers, where the down-scaled Vision Transformers not utilizing transfer learning exhibited notably lower performance as seen in Fig 3, but not necessary for the ResNet, due to being a convolutional neural network and inherently possesses and inductive bias for image classification.

The experiments on a fully balanced, non-treated dataset, as detailed S1 Appendix, indicated that classifying *P. aeruginosa* was more challenging. However, this was only apparent when using smaller models and during the single-frame subsampling experiments. This trend was not observed when using ViT/B-16, larger pretrained ResNets, or when performing spatially subsampled video classification.

All our training and model evaluation relies on defining the true class based on fluorescent staining of the final frame. Although the microfluidic device aims to study the lineage from one mother cell, the traps may occasionally have contained multiple species during growth prior to fluorescent capture. Some classification errors may therefore be due to artifacts, such as multiple species being present in the trap in the first frame, with one subsequently pushed out. Due to pressure variations, bacteria may also be sucked out of the trap and replaced by another species, or the trap may be empty at the start, with cells being pushed in during the experiment. Furthermore, poor fluorophore absorption by a species may indicate that a trap only contained one species when viewing the fluorescent images, where it, in reality, contained two. Manual inspection of the misclassified traps from the testing samples indicated that the aforementioned had occurred in six out of twelve misclassifications, see S1 Fig - S12 Fig. Another phenomenon that can occur is the appearance alteration of the bacteria image data, such as size, morphology, and structural irregularities, due to traps treated by antibiotics. This transformation occurs gradually, often becoming discernible in later frames rather than the initial ones. Despite these artifacts, the overall accuracy did not notably change when testing on later frames in the time-lapse, as seen in 4. Only a slight degradation is evident in the final few frames.

In the time-lapse experiments, accuracy improved with the number of frames available to the classifier, and the results slightly indicate that the order of the frames may be important and that temporal information from the time-lapses can be extracted to aid the network. In a future scenario in a clinical setting, accuracy may increase over time as video data is continuously acquired from the traps, providing increased confidence in the species classification.

A limitation of our method is its inability to handle mixed species within a single trap, however, it is expected to be a rare event in a clinical setting. Another obvious limitation in this proof-of-principle study is that we only extend our study to four species. More species and different isolates of the same species encountered in a clinical situation need to be addressed in future work.

We envision future applications where diagnostic tools containing microfluidic chips can quickly determine bacterial species and guide efficient treatment.

## 5 Materials and Methods

### 5.1 Bacterial strains, sample preparations, and antibiotics

In this study, we investigated four bacterial strains, representing both gram-negative and gram-positive cells, namely, *E. coli* K12 MG1655 (DA4201), *K. pneumoniae* (ATCC 13883), *P. aeruginosa* (DA6215) and *E. faecalis* (ATCC 51299). Glycerol stocks of each strain were independently cultured overnight in 5 ml Muller-Hinton (MH) medium at 37°C, then diluted 1:1000 times in a fresh MH medium supplemented with pluronic and allowed to grow for 2 hours at 37°C in a shaking incubator with a speed of 200 rpm. Subsequently, the strains were mixed in equal concentrations and loaded in a microfluidic chip. For susceptibility testing, the bacterial strains were treated with different antibiotics such as amoxicillin-clavulanate, ampicillin, ciprofloxacin, doripenem, nitrofurantoin, and gentamicin. The concentrations used correspond to MIC values for the *E. coli* suggested by the European Committee on Antimicrobial Susceptibility Testing (EUCAST).

### 5.2 Experimental setup and imaging conditions

We used the same experimental setup as in Kandavalli et al. 2022 [24], and Baltekin et al. 2017 [25]. Briefly, after loading the cells in a microfluidic chip, cells were exposed to media with and without antibiotics. For imaging, we used the Nikon Ti2-E inverted microscope equipped with a Plan Apo Lambda 100x oil immersion objective (Nikon). To monitor the growth of the bacteria, we captured phase contrast images every two minutes for an hour. After time-lapse imaging, the bacteria species in each trap in the microfluidic chip were identified as described in the genotyping protocol by Kandavalli et al. 2022. Next, we capture fluorescence images for each probe in four different fluorescence channels (Alexa Fluor 488, Cy3, Cy5, and Texas Red) corresponding to *E. faecalis*, *E. coli*, *K. pneumonia*, and *P. aeruginosa*, respectively.

### 5.3 Data selection

The microfluidic chips used in the experiments contained 34-44 traps at each capture position and were imaged at 162-180 capture positions depending on the experiment. Binary barcode labels were evenly laid out so that a unique identifier address could be assigned to each trap. Each phase-contrast and fluorescence microscopy frame was imaged with an image size of 1824 x 3888 pixels at 33 pixels/µm resolution.

To gather the data for this study, we developed custom image processing software to find and crop each growth trap and barcode label from the microscopy output. The phase-contrast stack was stabilized using rigid body transformations [26] using only the barcode labels as reference. Additionally, our image processing pipeline contained a procedure to shift the position of the fluorescence images to align them with the stabilized phase-contrast time-lapse stack. This procedure used a vertical projection of the pixel intensities of the different images into a one-dimensional vector, generating peaks at the horizontal location of the traps. The peaks from the fluorescence images and the phase-contrast projections were then shifted until alignment, and this shift was then replicated in the original images. The cropped images from the microfluidic chip’s traps measured between 50-54 pixels in width and 1400-1500 pixels in height. The cropping software and code for reproducing the deep-learning experiments are freely released alongside the dataset to facilitate further research [27].

Traps containing only one species in the mixed species experiments were manually selected by inspecting the final fluorescence image. This task was performed in a “semi-automated” way by sorting the traps by aggregated statistics of fluorescence pixel intensity values and visualizing the fluorescence channels side by side, noting the traps with only one bacteria species. A total of 3396 traps with corresponding time-lapses imaging single-species bacterial growth were extracted from seven different experiments divided into 684 *E. faecalis*, 770 *E. coli*, 1280 *K. pneumoniae*, and 662 *P. aeruginosa*. We maximized data extraction from these experiments, excluding only traps that were empty or contained multiple species as indicated by fluorescence signals. A number of train/test splits were created using predetermined random seeds, withholding 15% of the samples for testing. The dataset contained traps with and without antibiotic treatment during growth, as seen in Table 3.

The reason for using mixed species across several experiments was to mitigate the possibility of overfitting to experimental settings such as microscopy configurations or background features arising due to chip-to-chip variations, using so-called “Clever Hans”-prediction [28], or overinterpretation and adapting to non-salient features [29].

### 5.4 Single frame classification

When evaluating network performance on single frames, image classifiers were trained using only a single frame as input. In our primary experiments, the networks could only access the first frame in the time-lapses at test-time, as it holds greater clinical relevance to determine the correct species of the bacteria as early as possible. However, we also conducted experiments testing on subsequently later frames comparing the performance of ResNet-26 and ViT-B/16. During the training phase, all frames from the time-lapse were utilized, randomly selecting a frame for each sample in a mini-batch.

We compared the results of multiple ResNet architectures with the novel ViTs, both with and without transfer learning. While several architectural enhancements have been made to the ViT and ResNet (and ConvNet) architectures, our experimentation focused on their original versions to explore the inherent capabilities of each architecture. Furthermore, these foundational versions are inherently simpler, while newer architectures often incorporate networks with more complex operations and intricate pathways. Additionally, most new architectures are predominantly fine-tuned for natural images, with Imagenet commonly serving as the benchmark. Their performance on microscopic images remains largely uncharted. Recent studies indicate that aspects like training methods, augmentation techniques, and hyperparameter choices can significantly influence performance more than the actual model architecture [30].

#### 5.4.1 ResNets - Convolutional Neural Networks

In this study, we evaluated 21 different ResNet [8] architectures; the regular ImageNet ResNet with 18, 26, 34, 162, and 200 layers, and the more compact CIFAR-10 ResNet with 8, 14, 20, and 32 layers. Furthermore, we conducted ablation experiments with the CIFAR ResNet-8, reducing the base feature channel depth in the first layer to *r*, *r* = *{*3*, …,* 16*}*, named ResNet-8-r. Adopting this scaling strategy implied that we used feature channel depths *r*, 2*r*, and 4*r* in the network respectively. ResNet-8-16 is equivalent to CIFAR ResNet-8 since 16 is the default channel depth after the first layer.

The reason for using the smaller CIFAR ResNet variant was twofold; firstly, to determine whether this is a trivial problem or if robust gains can be obtained by using a deeper model with more parameters, secondly to assess the limitations if the models were to be employed in a diagnostics tool with limited compute resources.

The default ResNet variant used in our experiments was aimed at ImageNet-classification [4] in the original paper by He et al. [8]. It employed an initial set of 64 7×7 convolutional filters and max-pooling downsampling followed by a series of layers using 3×3 filters, grouped in pairs, referred to as “blocks.” Convolutions with a stride of 2 in the initial layer of an evenly spaced subset of these blocks were used, doubling feature-channel depth and halving spatial dimensions. Additive skip connections were introduced around each block. In instances where there was a discrepancy in channel depth, it was corrected by using 1×1 convolutions, transforming the feature maps to the same depth. Bottleneck blocks were used in place of the standard two-layer blocks for the deeper networks, allowing for less computationally expensive processing of the feature maps. The bottleneck blocks consisted of three layers; 1×1 filters reducing the channel depth, 3×3 filters processing at the lower channel depth, and finally, 1×1 filters transforming the feature map to the original dimension before the depth reduction.

The CIFAR10 ResNet variant used in our experiments was presented in the original paper by He et al. [8] and aimed to classify the smaller CIFAR10 dataset, consisting of 32×32 color images. It utilized a single 3×3 convolutional layer followed by the standard blocks but increased feature map depths to 16, 32, and 64, respectively, in the first layer at evenly spaced blocks. All ResNet variants included a global spatial average pooling layer, a fully connected layer, and a softmax activation function as a final layer. Batch normalization [7] was used before each activation.

#### 5.4.2 ViT - Vision Transformers

We evaluated 25 variants of the ViT architectures, the regular ViT-B with pretrained and randomly initialized weights using patch sizes 8 and 16, and downscaled versions of ViT-B/8. The scaling was performed as follows; the number of heads and the embedding dimension was linearly scaled down, keeping the number of layers constant at 12, analogously to the architectures presented in DeiT [31], ending up using single-head attention and an embedding dimension of 64. This scaling strategy was followed by proportionally scaling down the number of layers and embedding dimension, ending up in an embedding dimension of 16 and a depth of 3 layers, reducing the model complexity and observing the average accuracy. We name these architectures ViT-[*d_embed_*]-[*h*]-[*n_depth_*], where ViT-B/8 being equivalent with ViT-768-12-12.

In the ViT architecture [9], the input image was first split into *N* patches with a predefined size, and the patches were then flattened and linearly embedded into tokens with dimensionality *d_embed_*. A learned positional encoding was added to each token based on its position in the sequence. Each of the tokens was then projected into queries, keys, and values with dimensionality *d_q_*, *d_k_*, and *d_v_* using a set of projection matrices with learnable weights 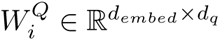, 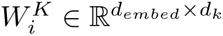, and 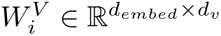. The projection was executed several times in parallel, called “heads” *h*, *∀i ∈ {*1*, …, h}*. The queries, keys, and values were then packed as rows into matrices *Q_i_*, *K_i_*, and *V_i_*, followed by an operation referred to as the self-attention mechanism.

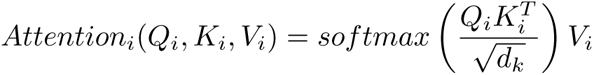

Where 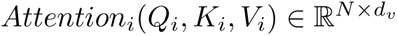. The outputs from all of the *h* heads were then concatenated column-wise into a matrix 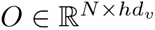 and then transformed back to the original dimensionality by a learnable matrix 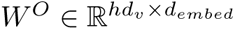.

Dimensionalities *d_q_*= *d_k_*= *d_v_*= *d_embed_/h* were chosen for the projection matrices to make the computational cost of the multi-head attention similar to single-head attention with full dimensionality. Additive skip-connections were added to this multi-head attention computation’s output, as well as a layer-wise normalization across all embeddings. A fully connected 2-layer feed-forward neural network then processed each embedding, with 4*d_embed_* hidden units, containing a skip connection and a layer-wise normalization. These computations were grouped into a block that was repeated *n_depth_*times. A learnable “classification token” was prepended to the patch embeddings and fed simultaneously through the network. The final output was based on this token being fed into a classification head consisting of a multilayer perceptron using a single layer at fine-tuning time.

#### 5.4.3 Decreasing resolution

Finally, the performance of ResNets and ViTs were compared when operating under decreased resolution. To approximately simulate using lower magnification microscopy, the Lanczos [32] interpolation was used for downsampling, which approximates the Sinc function. A convolution by a Sinc is equivalent to a low-pass filter in the frequency domain, removing high-frequency parts, thus effectively reducing the resolution. The image data was resized to a width of *w*, *w ∈ {*3, 4, 5, 6, 7, 8, 9, 10, 11, 12, 13, 16, 20, 24, 28, 32, 36, 40, 44, 48, 52*}*. A total of 21 pretrained ResNet-26 networks and 16 ViT-B/8 networks were trained using the subsampled image data at both train and test time.

### 5.5 Time-lapse classification and temporal features

To investigate whether temporal dynamics were essential for classification accuracy, several “R(2+1)D”-Video ResNet classification networks introduced by Tran et al. [11] were trained on the time-lapse data. A total of 32 frames were obtained from each trap, duplicating the last frame when the number of frames was lower than 32 in the time-lapse and cropping the time-lapse longer than 32 frames to 32 frames. In total, 32 networks were trained using different numbers of frames as input, *n_frames_ ∈ {*1, 2*, …,* 32*}*. Next, identical video classification networks were trained but randomly shuffling the frames before input. The networks trained on time-lapses with fewer than 32 frames utilized temporal jittering, [11] randomly sampling *n_frames_* consecutive frames from the time-lapse during training. Testing was performed starting from the first frame in the time-lapse. We hypothesize that if temporal information mattered, the networks processing shuffled frames would consistently yield a lower accuracy compared to the networks trained on frames in the correct order.

Furthermore, we conducted full-frame (32 frames) video classification experiments using spatially sub-sampled data, analogous to the single-frame subsampling.

Due to the surprisingly high accuracy achieved at extremely low resolutions, we conducted an experiment where the pixels were shuffled during the inference stage to investigate whether the network’s decision was based solely on image intensity for the smallest image sizes. The network failed completely at this task with a class-average accuracy of 25% if the pixels in the video were shuffled, indicating that information such as mean and standard deviation of intensity is insufficient for classification of bacterial species and the network must learn spatiotemporal features, even when processing low-resolution images. Supporting Videos 1-4 show that subtle cell features and temporal dynamics associated with bacterial growth remain discernable even under heavy subsampling.

The R(2+1)D Video ResNet network used repeated blocks with 2D spatial convolutions, processing each frame separately, followed by 1D temporal convolutions, fusing temporal information across frames [11]. The networks were pretrained on the Kinetics-400 dataset [33] and fine-tuned on our bacterial dataset.

### 5.6 Training regimes

The deep learning models were all trained with stochastic gradient descent using the AdamW [6] method, applying standard augmentations such as random crops, horizontal flips, randomly shifting, scaling, and rotating, as well as randomly chaining the brightness and contrast of the image. A cosine learning rate scheduler with warm restarts [34] was used, using five warm-up epochs with a decay rate of 0.5. The cycles had 100 epochs each for the single-frame classification networks and 75 for the video classification networks.

R(2+1)D Video ResNet networks and single frame classifications were trained with batch sizes of 8 and 32, respectively. The learning rates were 0.0002 for Video ResNets, 0.001 for ResNets, and 0.0001 for ViTs. However, due to convergence issues, the 34-layer ResNet’s learning rate was reduced to 0.0001. The Video ResNets were trained for 150 epochs, single-frame ResNets for 300 epochs, and the ViT for 500 epochs. The ViTs required more epochs to converge, possibly attributed to the lower learning rate and not having an inductive bias for image classification. We employed a random weighted sampler to sample the data equally and address the minor class imbalance. After random cropping, each frame had a size of 52×382 pixels.

The models were trained using an Nvidia A100 40GB GPU, which was partitioned using the MIG (Multi-Instance GPU) 3g.20gb configuration, which approximately equates to halving the capabilities of the original GPU. Any GPU manufactured in the early 2020s and late 2010s should suffice to train the models. Training times were 1-2 hours for the ResNet models, 1-10 hours for the ViTs, and 1-12 hours for the Video ResNets. Inference time is highly device-specific but demonstrated subsecond speeds when using our system and is directly correlated to the number of Floating point operations presented in Tables 1 and 2.

**Table 2.** Model results: Video classification.

| Video Resnet "R(2+1)D" |  |  |  | Video Resnet "R(2+1)D" |  |  |  |
| --- | --- | --- | --- | --- | --- | --- | --- |
| Frames | Parameters | FLOPS | Accuracy | Resolution | Parameters | FLOPS | Accuracy |
| 1 | 31.30 M | 15.28 GFlops | 97.55 ± 0.35% | 3x22 | 31.30 M | 1.56 GFlops | 98.68 ± 0.24% |
| 2 | 31.30 M | 19.75 GFlops | 98.39 ± 0.75% | 4x29 | 31.30 M | 1.99 GFlops | 98.53 ± 0.34% |
| 3 | 31.30 M | 27.61 GFlops | 97.83 ± 0.54% | 5x37 | 31.30 M | 3.54 GFlops | 99.06 ± 0.41% |
| 4 | 31.30 M | 32.08 GFlops | 98.06 ± 0.48% | 6x44 | 31.30 M | 4.19 GFlops | 98.96 ± 0.27% |
| 5 | 31.30 M | 43.60 GFlops | 98.20 ± 0.42% | 7x51 | 31.30 M | 5.26 GFlops | 98.85 ± 0.28% |
| 6 | 31.30 M | 48.07 GFlops | 98.01 ± 0.34% | 8x59 | 31.30 M | 6.12 GFlops | 98.99 ± 0.19% |
| 7 | 31.30 M | 55.93 GFlops | 98.17 ± 0.58% | 9x66 | 31.30 M | 9.44 GFlops | 99.22 ± 0.35% |
| 8 | 31.30 M | 60.39 GFlops | 98.20 ± 0.70% | 10x73 | 31.30 M | 10.44 GFlops | 99.15 ± 0.19% |
| 9 | 31.30 M | 75.68 GFlops | 98.42 ± 0.61% | 11x81 | 31.30 M | 12.46 GFlops | 99.23 ± 0.16% |
| 10 | 31.30 M | 80.15 GFlops | 98.30 ± 0.31% | 12x88 | 31.30 M | 13.88 GFlops | 99.35 ± 0.33% |
| 11 | 31.30 M | 88.00 GFlops | 98.45 ± 0.35% | 13x96 | 31.30 M | 16.97 GFlops | 99.09 ± 0.17% |
| 12 | 31.30 M | 92.47 GFlops | 98.41 ± 0.50% | 16x118 | 31.30 M | 22.43 GFlops | 99.32 ± 0.43% |
| 13 | 31.30 M | 103.99 GFlops | 98.56 ± 0.50% | 20x147 | 31.30 M | 37.44 GFlops | 99.40 ± 0.34% |
| 14 | 31.30 M | 108.46 GFlops | 98.69 ± 0.30% | 24x176 | 31.30 M | 51.90 GFlops | 99.36 ± 0.47% |
| 15 | 31.30 M | 116.32 GFlops | 98.95 ± 0.48% | 28x206 | 31.30 M | 70.73 GFlops | 99.33 ± 0.24% |
| 16 | 31.30 M | 120.79 GFlops | 98.70 ± 0.41% | 32x235 | 31.30 M | 90.59 GFlops | 99.21 ± 0.44% |
| 17 | 31.30 M | 136.07 GFlops | 98.78 ± 0.42% | 36x264 | 31.30 M | 117.85 GFlops | 99.32 ± 0.46% |
| 18 | 31.30 M | 140.54 GFlops | 99.00 ± 0.33% | 40x294 | 31.30 M | 141.99 GFlops | 99.36 ± 0.32% |
| 19 | 31.30 M | 148.40 GFlops | 99.25 ± 0.43% | 44x323 | 31.30 M | 172.85 GFlops | 99.38 ± 0.33% |
| 20 | 31.30 M | 152.87 GFlops | 98.99 ± 0.38% | 48x353 | 31.30 M | 202.55 GFlops | 99.31 ± 0.51% |
| 21 | 31.30 M | 164.39 GFlops | 99.13 ± 0.32% | 52x382 | 31.30 M | 241.58 GFlops | 99.53 ± 0.40% |
| 22 | 31.30 M | 168.86 GFlops | 99.04 ± 0.30% | | | | |
| 23 | 31.30 M | 176.71 GFlops | 99.01 ± 0.20% | | | | |
| 24 | 31.30 M | 181.18 GFlops | 99.38 ± 0.16% | | | | |
| 25 | 31.30 M | 196.47 GFlops | 99.35 ± 0.24% | | | | |
| 26 | 31.30 M | 200.94 GFlops | 99.21 ± 0.26% | | | | |
| 27 | 31.30 M | 208.79 GFlops | 99.55 ± 0.25% | | | | |
| 28 | 31.30 M | 213.26 GFlops | 99.51 ± 0.25% | | | | |
| 29 | 31.30 M | 224.78 GFlops | 99.48 ± 0.17% | | | | |
| 30 | 31.30 M | 229.25 GFlops | 99.45 ± 0.39% | | | | |
| 31 | 31.30 M | 237.11 GFlops | 99.47 ± 0.30% | | | | |
| 32 | 31.30 M | 241.58 GFlops | 99.32 ± 0.40% | | | | |

Average test accuracies and their corresponding standard deviations were computed across five retraining iterations, each utilizing a predetermined random seed.

**Table 3.**
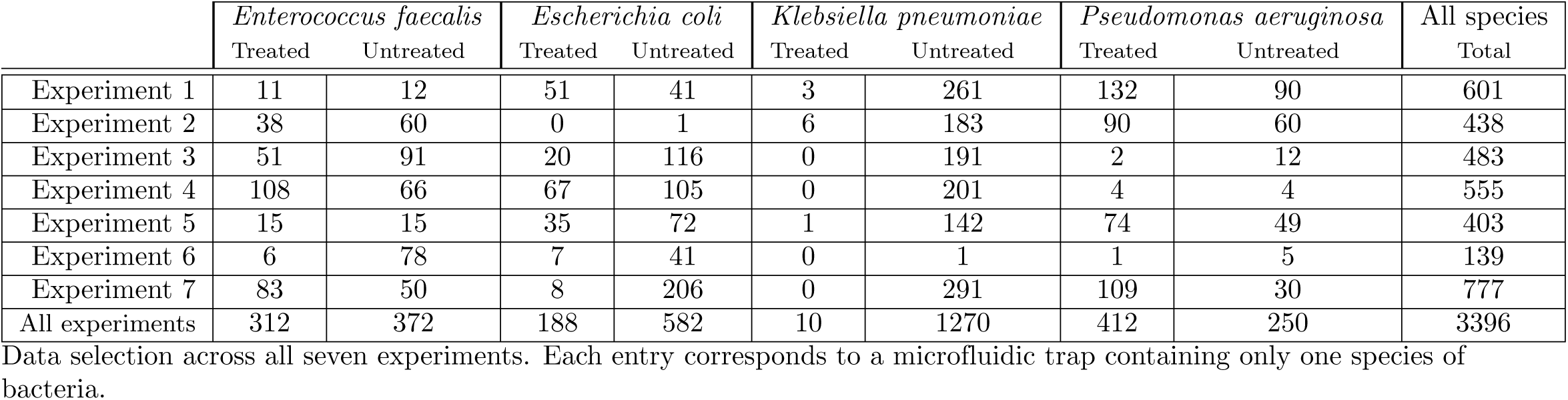
Data selection.

|  | *Enterococcus faecalis* |  | *Escherichia coli* |  | *Klebsiella pneumoniae* |  | *Pseudomonas aeruginosa* |  | All species |
| --- | --- | --- | --- | --- | --- | --- | --- | --- | --- |
|  | Treated | Untreated | Treated | Untreated | Treated | Untreated | Treated | Untreated | Total |
| Experiment 1 | 11 | 12 | 51 | 41 | 3 | 261 | 132 | 90 | 601 |
| Experiment 2 | 38 | 60 | 0 | 1 | 6 | 183 | 90 | 60 | 438 |
| Experiment 3 | 51 | 91 | 20 | 116 | 0 | 191 | 2 | 12 | 483 |
| Experiment 4 | 108 | 66 | 67 | 105 | 0 | 201 | 4 | 4 | 555 |
| Experiment 5 | 15 | 15 | 35 | 72 | 1 | 142 | 74 | 49 | 403 |
| Experiment 6 | 6 | 78 | 7 | 41 | 0 | 1 | 1 | 5 | 139 |
| Experiment 7 | 83 | 50 | 8 | 206 | 0 | 291 | 109 | 30 | 777 |
| All experiments | 312 | 372 | 188 | 582 | 10 | 1270 | 412 | 250 | 3396 |

Data selection across all seven experiments. Each entry corresponds to a microfluidic trap containing only one species of bacteria.

All networks were retrained five consecutive times with different predetermined random seeds, ensuring reproducibility and uniformity in the training and evaluation process, as they were all trained and assessed using identical train/test partition setups, augmentation sequences, and weight initializations. The mean and standard deviation of the average accuracies of the classifications were calculated and presented in the results section.

The networks were pretrained on 3-channel natural images, on the contrary, the phase-contrast image data had only a single channel as input. The pretrained filters in the first layer with filter size n x n x 3 were reshaped to n x n x 1, computing the average across the last axis to accommodate for this. The open-source libraries PyTorch [35] and PyTorch Image Models [36] were used, for full details refer to the released software and dataset [27].

## 6 Acknowledgments

We acknowledge funding from the Swedish Foundation for Strategic Research, SSF ARC19-0016. The computations were enabled by the Berzelius supercomputing resource provided by the National Supercomputer Centre at Linköping University and supported by the Knut and Alice Wallenberg Foundation. Additionally, computations were enabled by the Alvis cluster provided by the National Academic Infrastructure for Supercomputing in Sweden (NAISS) at Chalmers University of Technology, partially funded by the Swedish Research Council through grant agreement no. 2022-06725. We would also like to thank Anders Hast for valuable input on the manuscript. Johan Elf has founded and is partly engaged in the AMR diagnostics company Sysmex Astrego AB.

## 7 Supporting information

### 7.1 Misclassifications

The following supporting figures are misclassifications made by the models in Fig 9, fine-tuned on the bacteria dataset and tested from the first frame. Labels *N*, *N ∈ {*0*, …,* 35*}* show ordered phase-contrast image data for a single trap in the time-lapse, labels *L*, *L ∈ {A, B, C, D}* show the final fluorescent imaging after the last frame showing probes Alexa488, Cy3, Cy5, and Txr (Texas Red) respectively, corresponding to species *E. faecalis*, *E. coli*, *K. pneumoniae*, and *p. aeruginosa*. The single-channel grayscale image data is visualized using the Viridis color map.

**S1 Fig.**
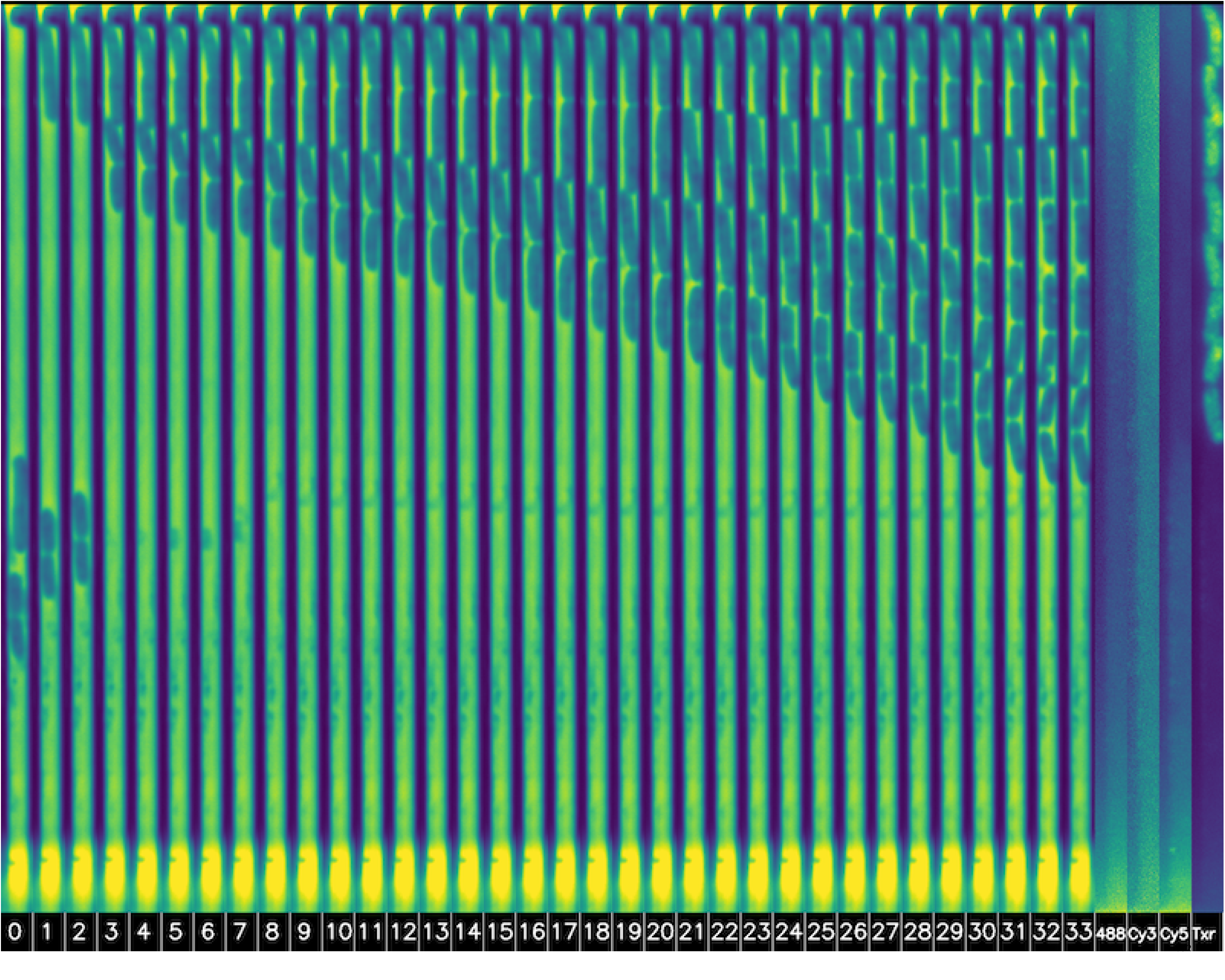
Misclassified by ResNet-26. True label *p. aeruginosa*, classified as *E. faecalis*. The ResNet may have confused the stop at the top of the trap as a coccus. The upper half of the trap was empty in the first frame.

**S2 Fig.**
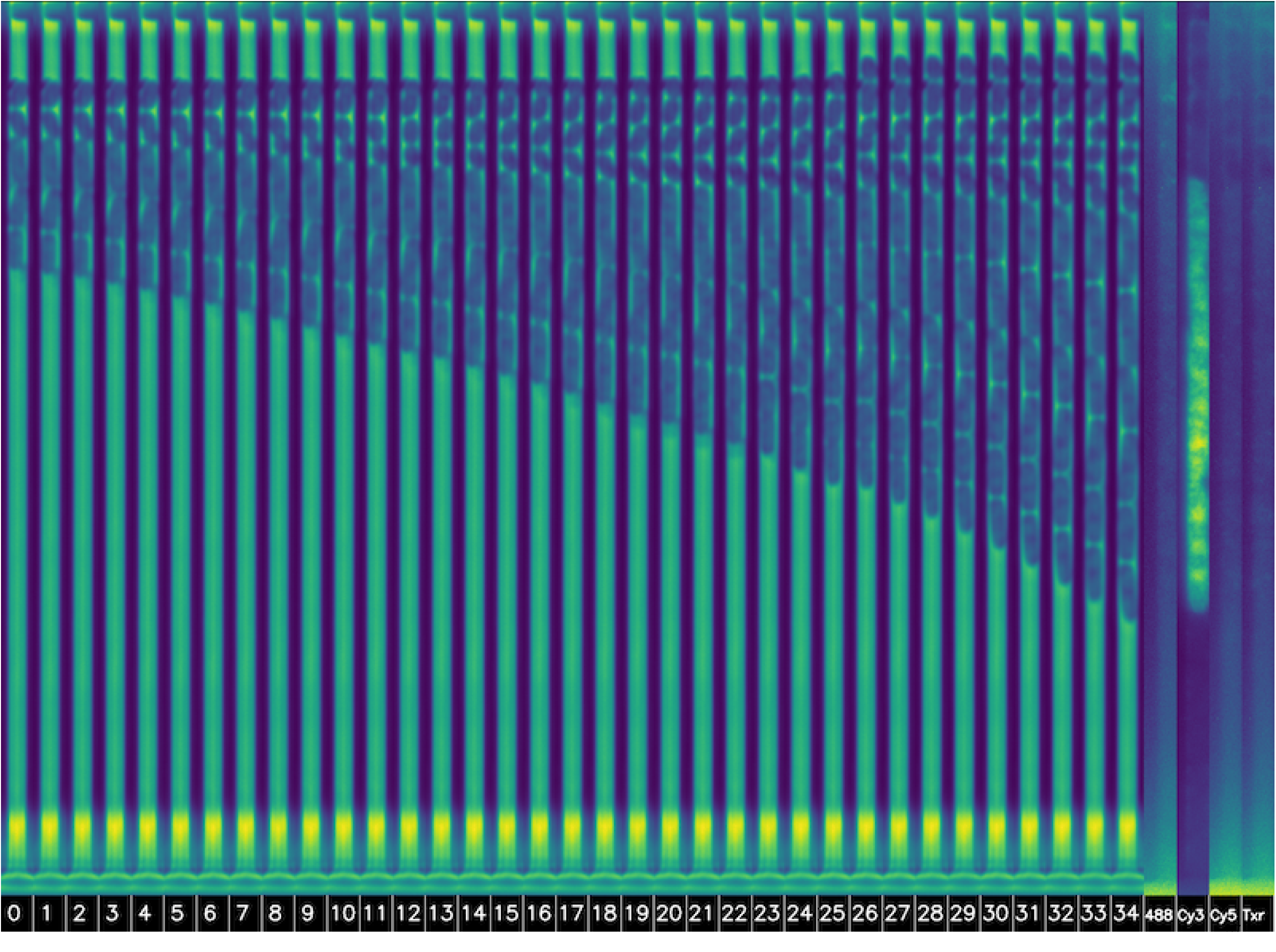
Misclassified by ResNet-26. True label *E. coli*, classified as *E. faecalis*. It appears to have been cocci in the trap that avoided staining, and the trap clearly did not only contain a single bacterial species.

**S3 Fig.**
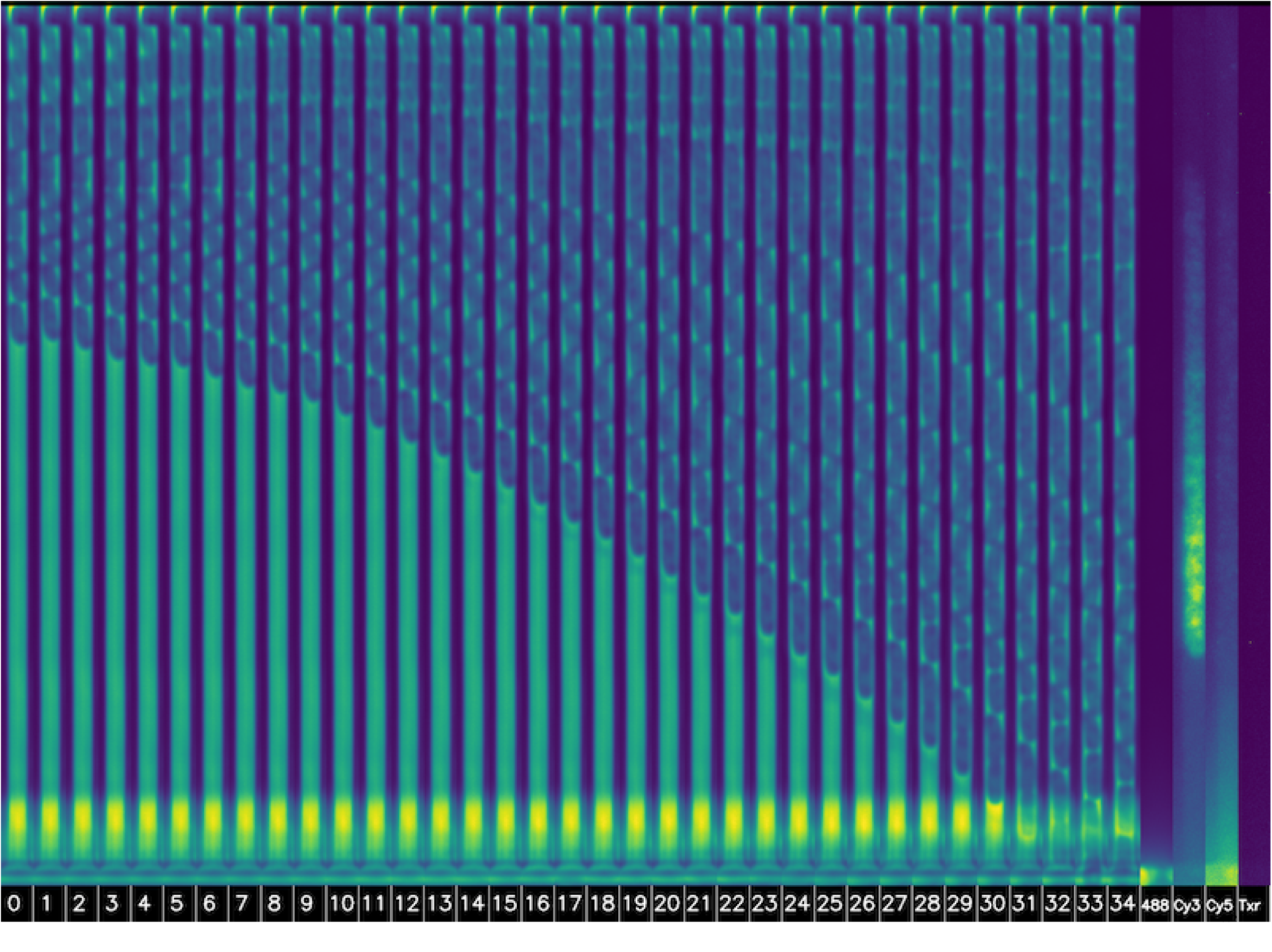
Misclassified by ResNet-26. *E. coli*, classified as p. aeruginosa. Both species are rods with similar shapes and are easily confused, and the fluorescent staining indicates that there may be two bacterial species in the trap, where one has avoided staining.

**S4 Fig.**
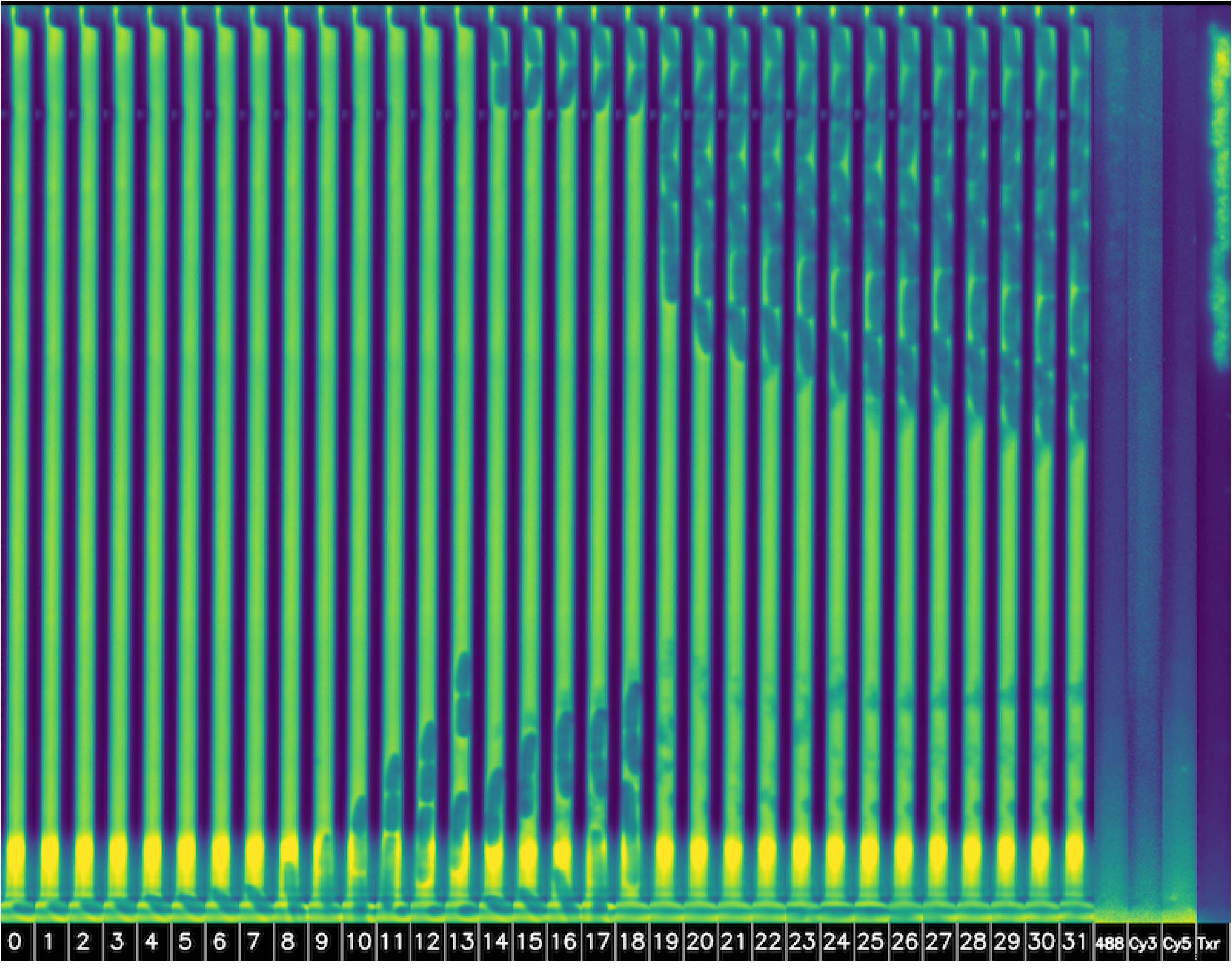
Misclassified by ResNet-26. True label *p. aeruginosa*, classified as *K. pneumoniae*. The trap was empty in the first frames.

**S5 Fig.**
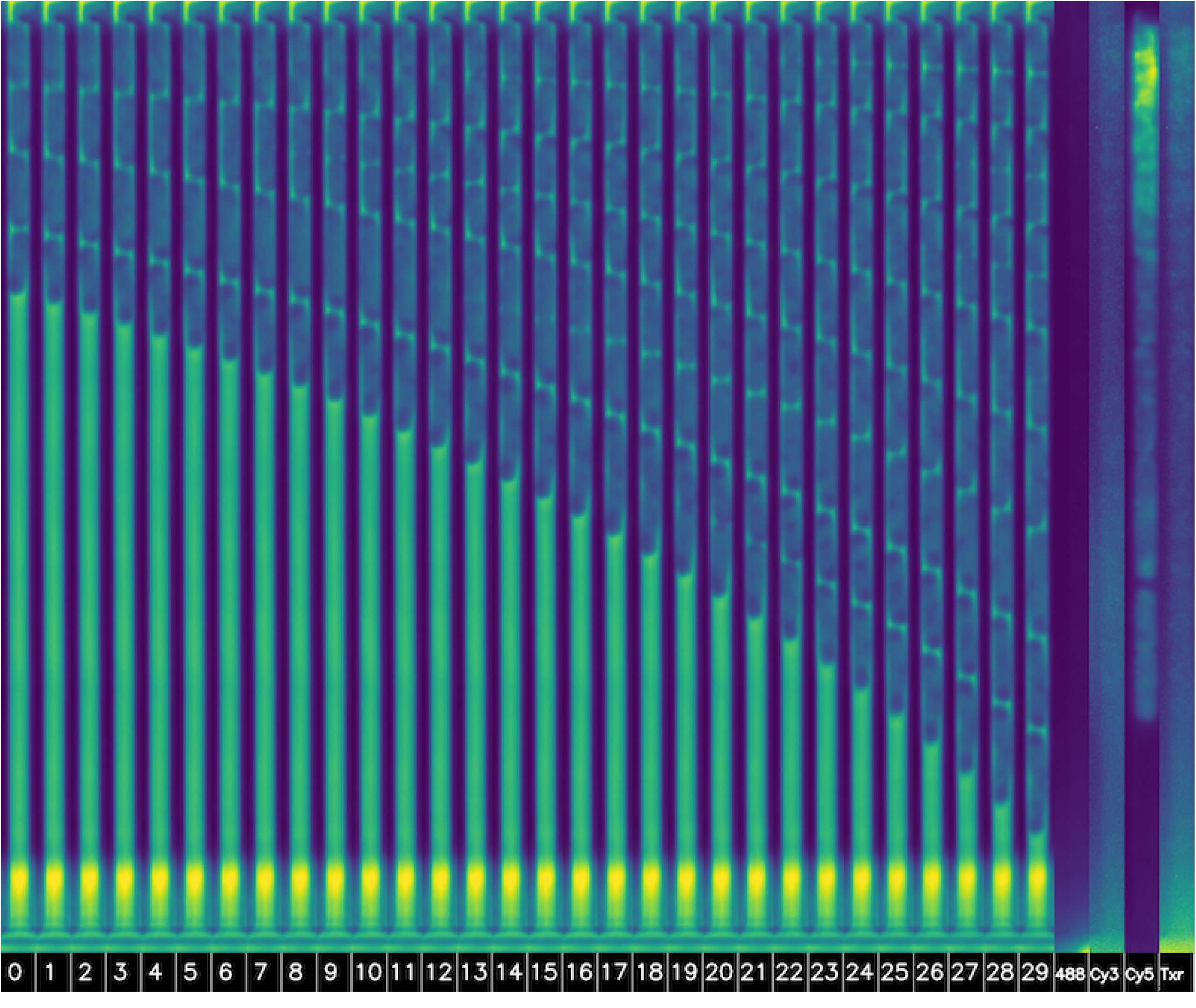
Misclassified by ResNet-26. True label *K. pneumoniae*, classified as *E. coli*. Both species are rods with similar shapes and are easily confused.

**S6 Fig.**
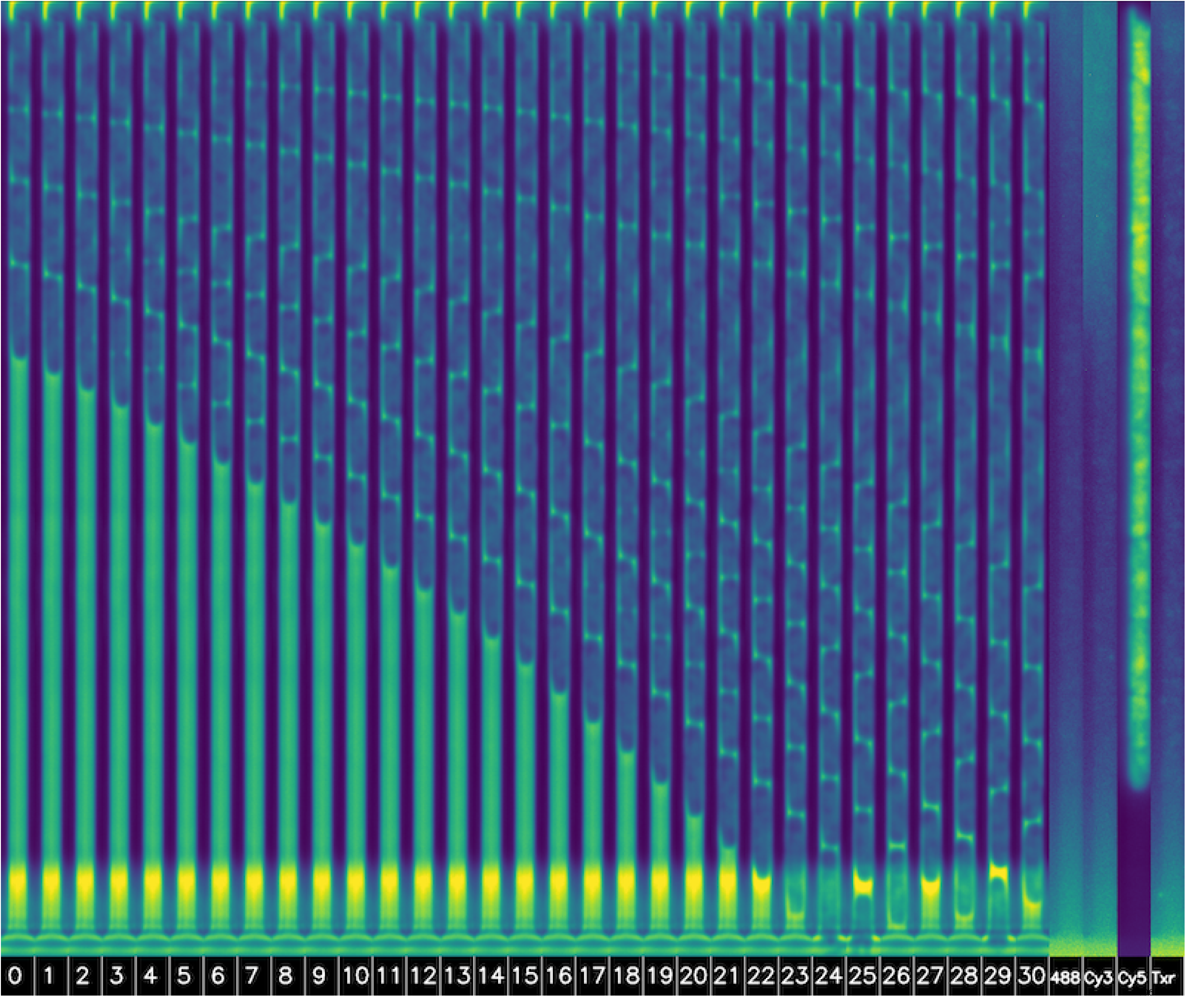
Misclassified by ResNet-26. True label *K. pneumoniae*, classified as *E. coli*. Both species are rods with similar shapes and are easily confused.

**S7 Fig.**
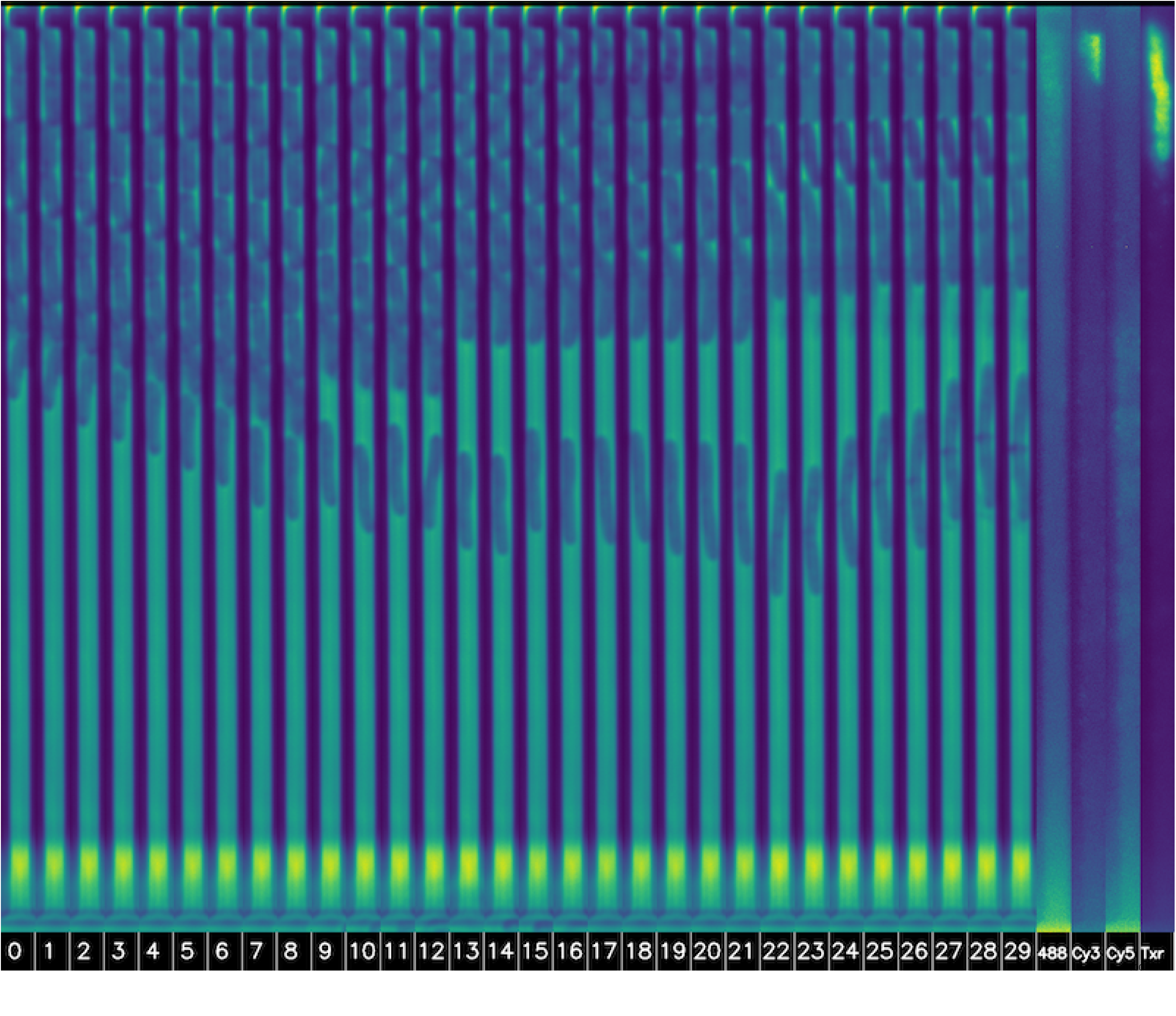
Misclassified by ResNet-26. True label *p. aeruginosa*, classified as *E. coli*. Both species are rods with similar shapes and are easily confused. It appears to have been several species in the trap that avoided staining under heavy antibiotic treatment.

**S8 Fig.**
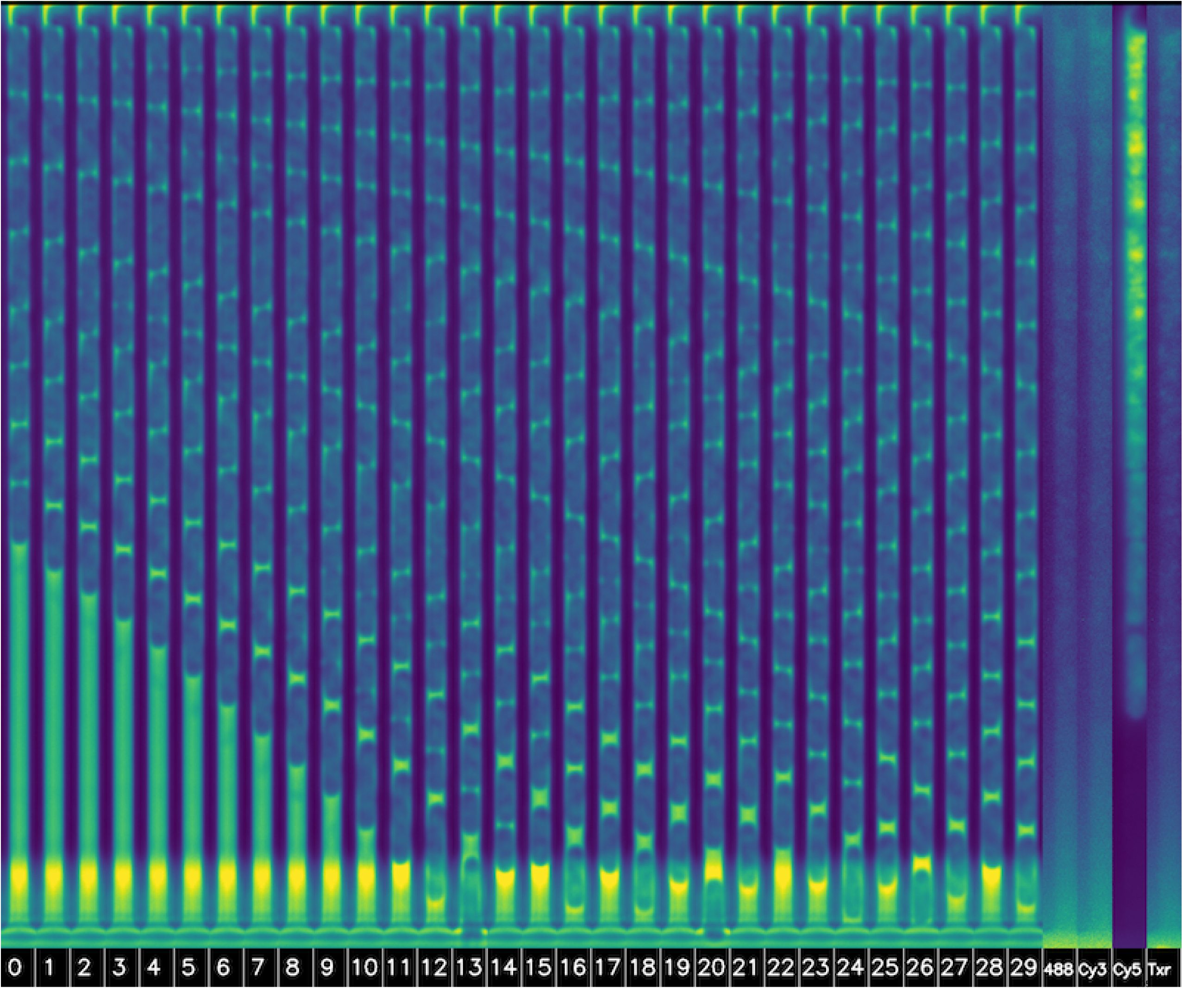
Misclassified by ResNet-26. True label *K. pneumoniae*, classified as *E. coli*. Both species are rods with similar shapes and are easily confused.

**S9 Fig.**
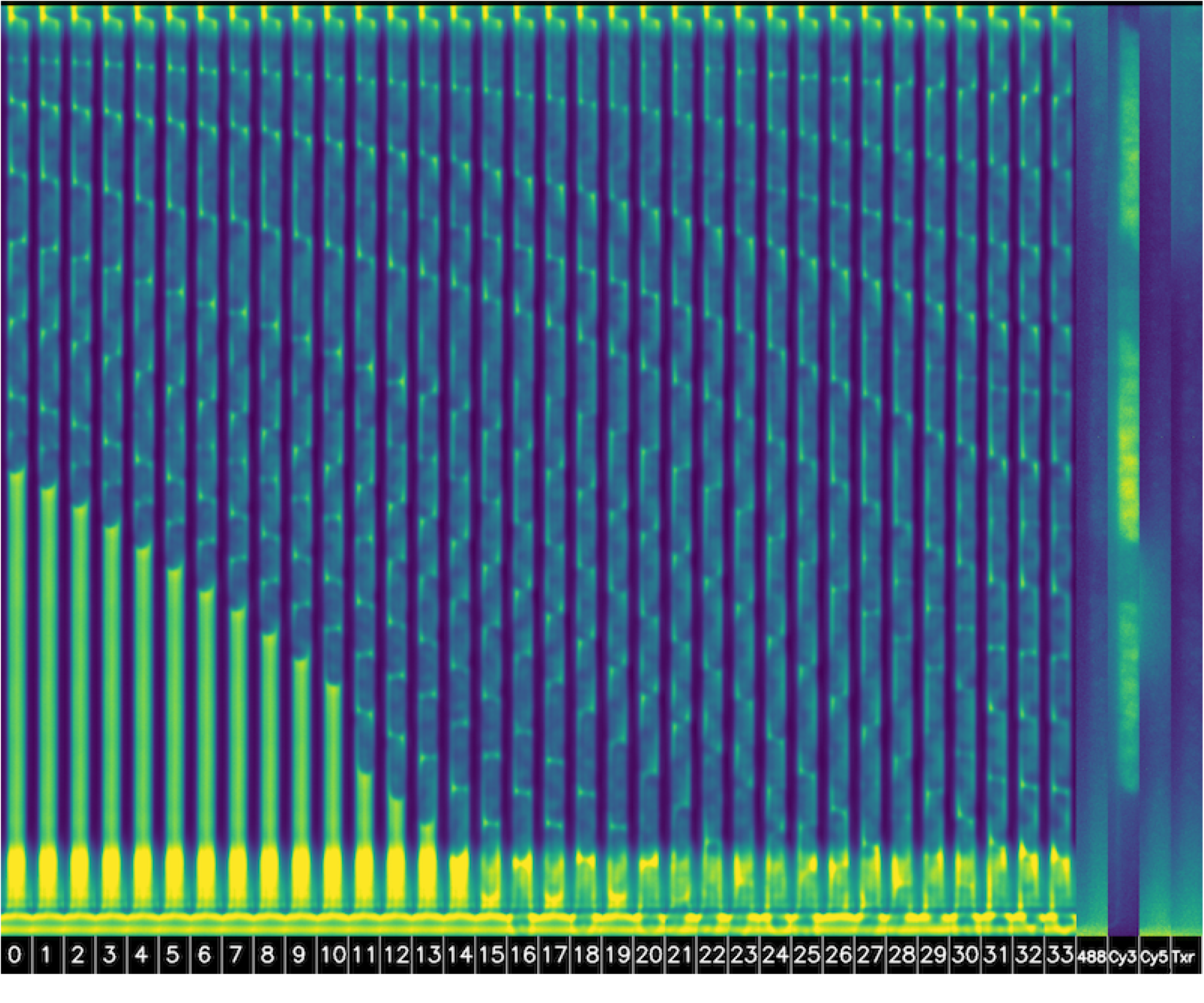
Misclassified by ViT/B. True label *E. coli*, classified as *K. pneumoniae*. Both species are rods with similar shapes and are easily confused.

**S10 Fig.**
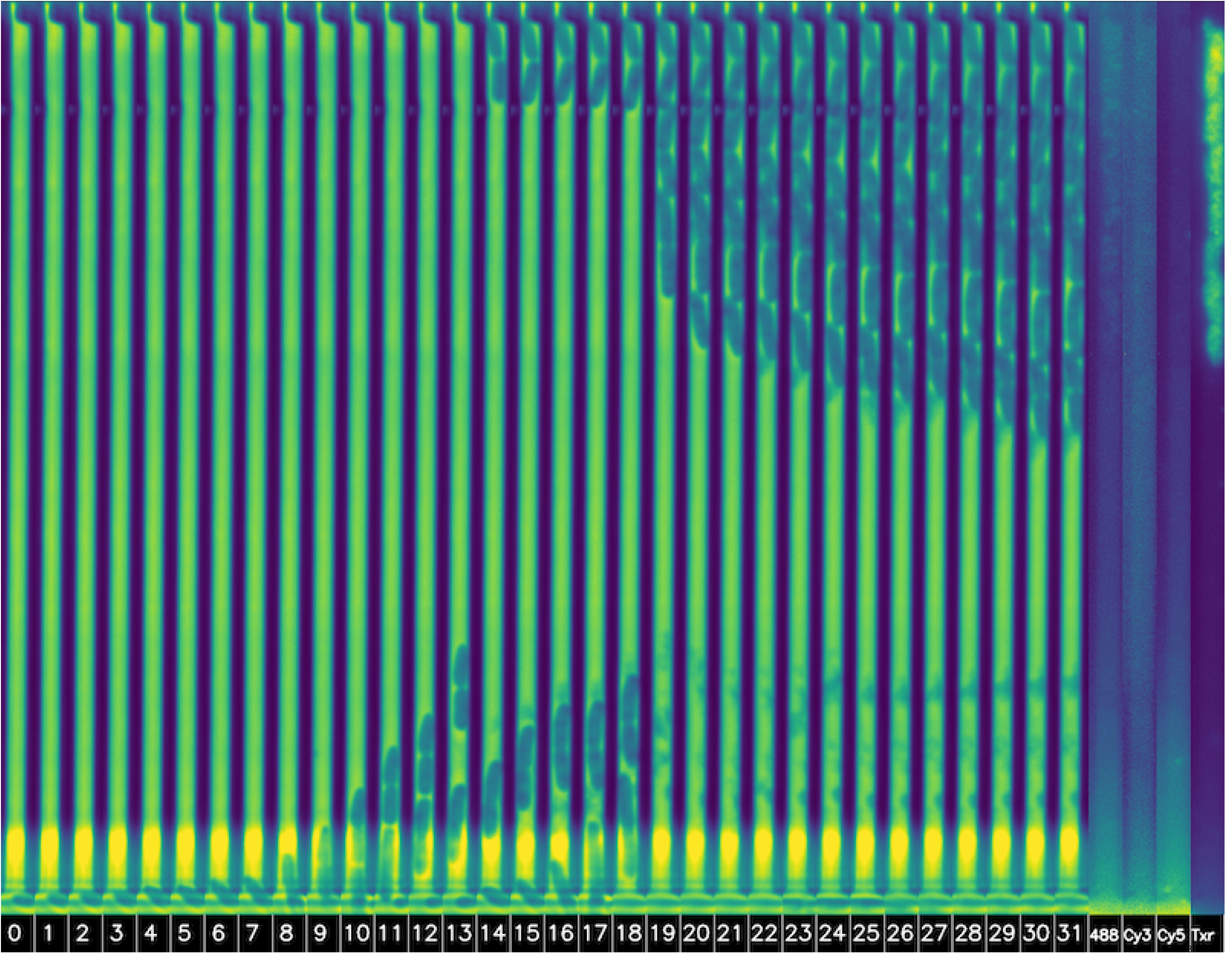
Misclassified by ViT/B. True label *p. aeruginosa*, classified as *E. faecalis*. The ViT possibly confused the stop at the top of the trap as a coccus. The trap was empty in the first frame.

**S11 Fig.**
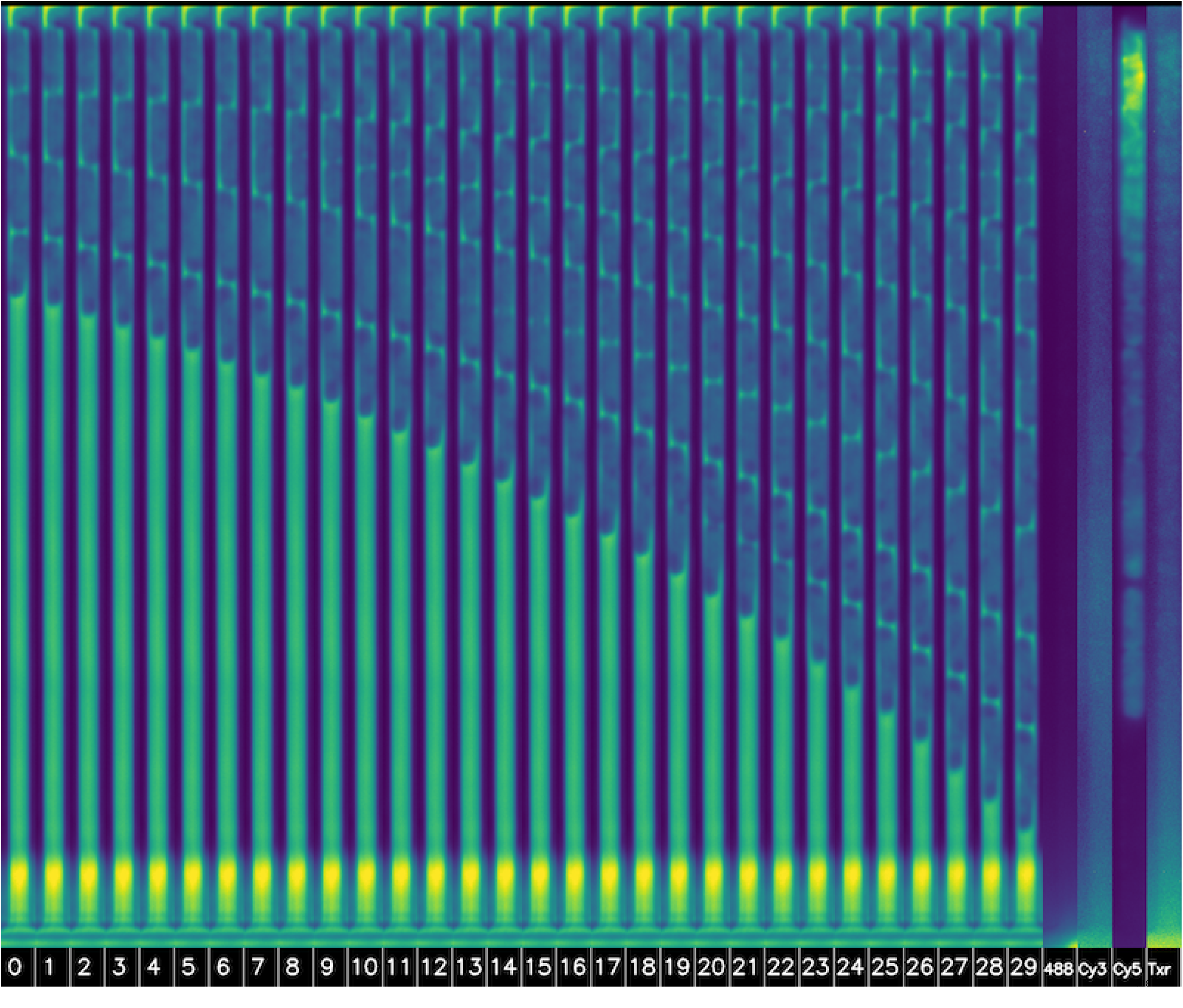
Misclassified by ViT/B. True label *K. pneumoniae*, classified as *E. coli*. Both species are rods with similar shapes and are easily confused.

**S12 Fig.**
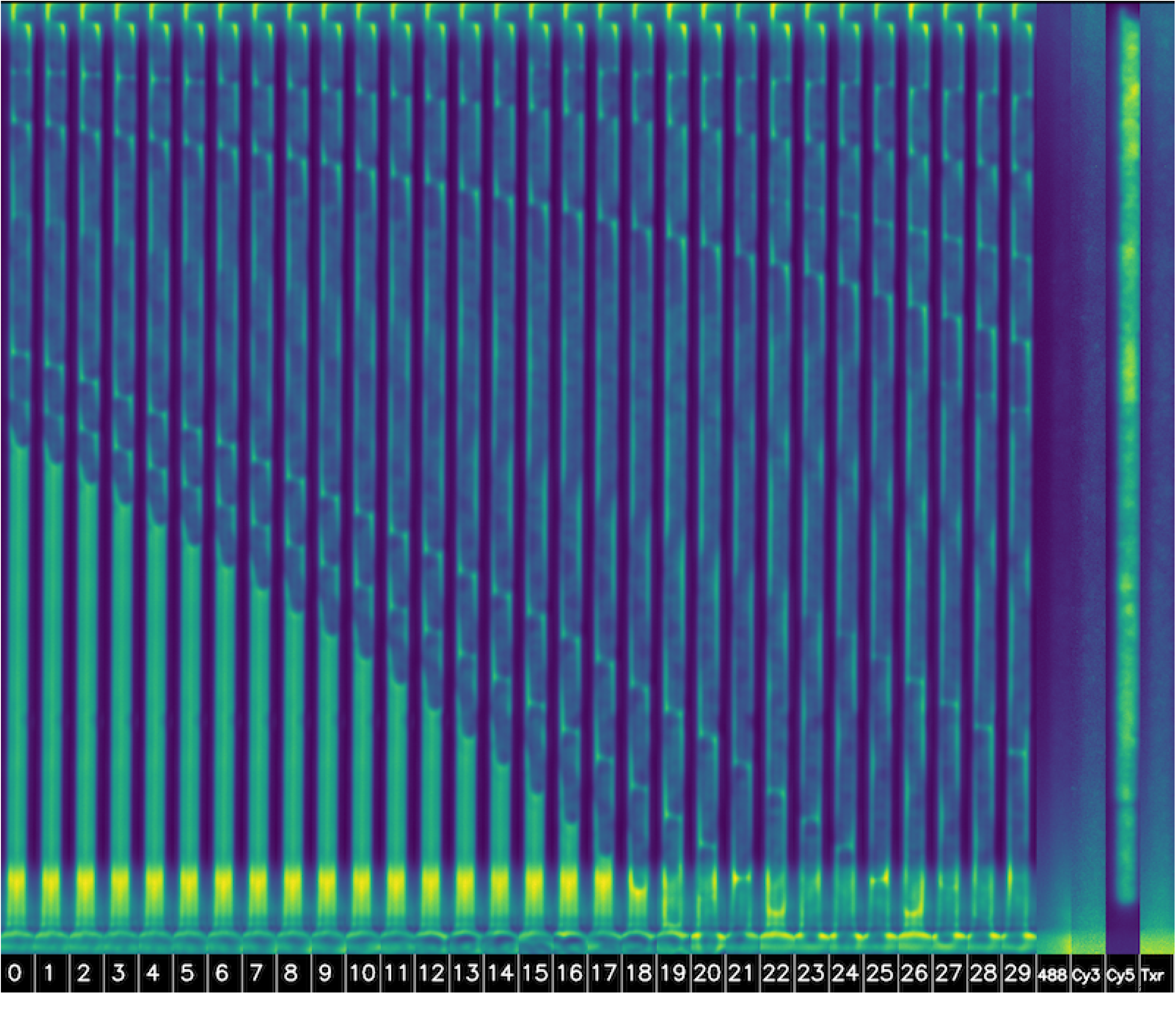
Misclassified by R(2+1)D. True label *K. pneumoniae*, classified as *E. coli*. Both species are rods with similar shapes and are easily confused.

### 7.2 Video time-lapses

The following supporting videos represent the data that is inputted into the neural networks, with one sample per species. All subsampling steps are displayed side by side for the same sample, ranging from the heavily subsampled image size of 3×22 on the left to the original 52×382 on the right. On the very left, the pixels are shuffled for the lowest resolution in our experiments. The single-channel grayscale video data is visualized using the Viridis color map.

**S1 Video Time-lapse video of *E. faecalis* reproducing in a trap**

**S2 Video Time-lapse video of *E. coli* reproducing in a trap**

**S3 Video Time-lapse video of *K. pneumoniae* reproducing in a trap**

**S4 Video Time-lapse video of *p. aeruginosa* reproducing in a trap**

### 7.3 Appendices

**S1 Appendix Fully balanced experiments using non-treated traps** A balanced dataset of 250 time-lapses per species was extracted from the untreated samples, trained, and tested on downscaled versions of CIFAIR-10 ResNet and ViT-B/8.

