## Supporting Appendix 1 for "Label-free deep learning-based species classification of bacteria imaged by phase-contrast microscopy"

### S1 Appendix: Fully balanced experiments using non-treated traps

The results showed a consistently low classification accuracy of *Pseudomonas aeruginosa* in the single-frame experiments, especially when using smaller models and subsampled low-resolution images. It was suspected that this could be because this particular species contained only 250 out of 662 untreated time-lapses. In response, we conducted additional experiments, selecting only 250 untreated traps from each species, using a 15% train/test split as before, retraining the downscaled versions of ResNet-8 and ViT-B/8, testing on the first frame. The results showed similar tendencies on this minimal test set. *Pseudomonas aeruginosa* attained the lowest accuracy in 20 and the second lowest in 6 of the models, as shown in Tables 1 and 2.

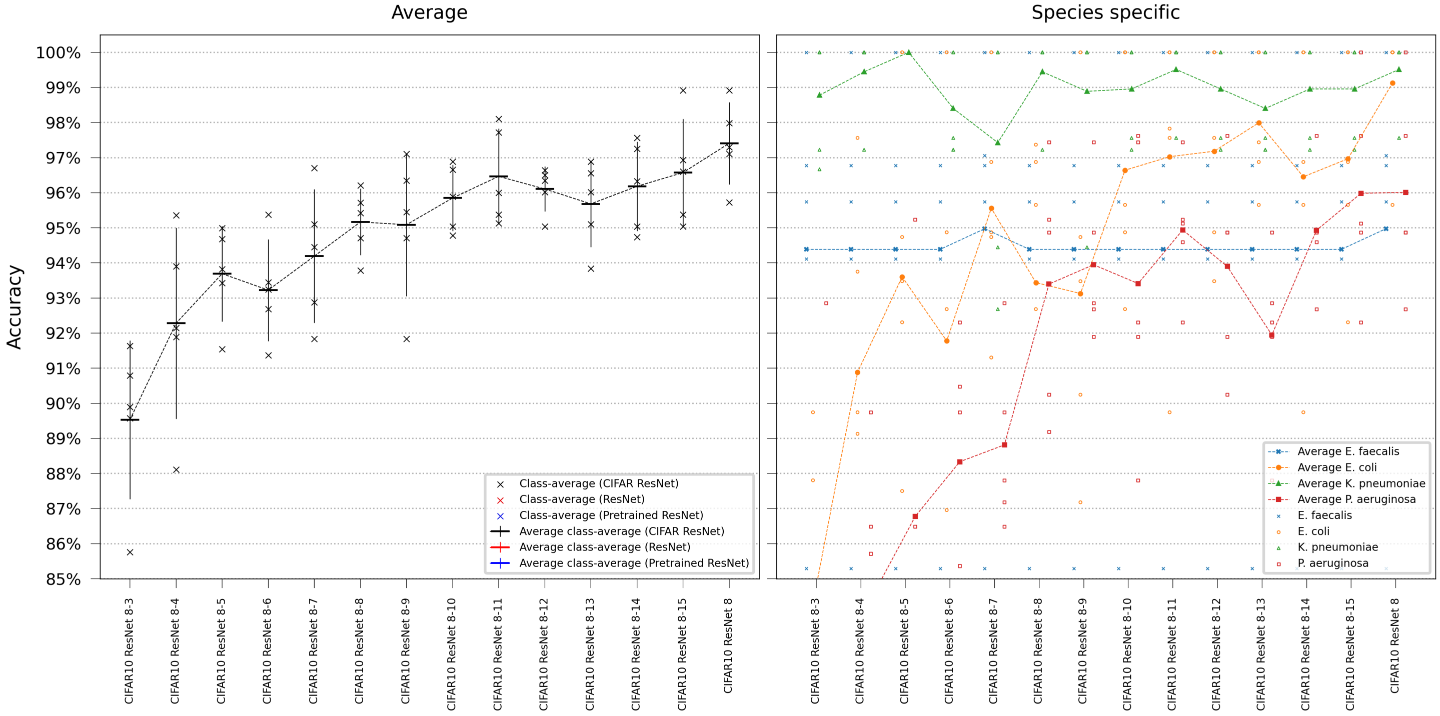


Figure 1: Model comparison performing single-frame classification of the first frame in the time-lapse using downscaled versions of ResNet-8. Error bars represent the standard deviation in class-average accuracy from the five retrainings. Scatter plots depict the class-specific accuracy of all individual classifiers and the average class-specific accuracy over the five retrainings for each model. To reduce overplotting, a minor jitter was introduced along the categorical axis of the species-specific scatter plot. Lines are included not for interpolation or statistical inference purposes but to visually guide readers in tracking mean values on the ordinal scale.

| Model name | Enerococus facuialis | E coli | Klebsiella | Pseudamonas |
|---|---|---|---|---|
| CIFAR10 ResNet 8-3 | $94.39 \pm 5.52\%$ | $84.31 \pm 5.67\%$ | $98.78 \pm 1.69\%$ | $80.65 \pm 8.27\%$ |
| CIFAR10 ResNet 8-4 | $94.39 \pm 5.52\%$ | $90.88 \pm 5.04\%$ | $99.44 \pm 1.24\%$ | $84.41 \pm 4.49\%$ |
| CIFAR10 ResNet 8-5 | $94.39 \pm 5.52\%$ | $93.60 \pm 4.50\%$ | $100.00 \pm 0.00\%$ | $86.78 \pm 4.89\%$ |
| CIFAR10 ResNet 8-6 | $94.39 \pm 5.52\%$ | $91.78 \pm 6.25\%$ | $98.40 \pm 1.47\%$ | $88.34 \pm 3.60\%$ |
| CIFAR10 ResNet 8-7 | $94.97 \pm 5.64\%$ | $95.56 \pm 3.19\%$ | $97.43 \pm 3.58\%$ | $88.81 \pm 2.57\%$ |
| CIFAR10 ResNet 8-8 | $94.39 \pm 5.52\%$ | $93.44 \pm 5.26\%$ | $99.44 \pm 1.24\%$ | $93.40 \pm 3.52\%$ |
| CIFAR10 ResNet 8-9 | $94.39 \pm 5.52\%$ | $93.13 \pm 4.84\%$ | $98.89 \pm 2.48\%$ | $93.95 \pm 2.24\%$ |
| CIFAR10 ResNet 8-10 | $94.39 \pm 5.52\%$ | $96.64 \pm 3.25\%$ | $98.96 \pm 1.43\%$ | $93.41 \pm 4.15\%$ |
| CIFAR10 ResNet 8-11 | $94.39 \pm 5.52\%$ | $97.03 \pm 4.23\%$ | $99.51 \pm 1.09\%$ | $94.94 \pm 1.83\%$ |
| CIFAR10 ResNet 8-12 | $94.39 \pm 5.52\%$ | $97.18 \pm 2.96\%$ | $98.96 \pm 1.43\%$ | $93.90 \pm 2.88\%$ |
| CIFAR10 ResNet 8-13 | $94.39 \pm 5.52\%$ | $97.99 \pm 1.94\%$ | $98.40 \pm 1.47\%$ | $91.95 \pm 2.58\%$ |
| CIFAR10 ResNet 8-14 | $94.39 \pm 5.52\%$ | $96.45 \pm 4.21\%$ | $98.96 \pm 1.43\%$ | $94.93 \pm 1.76\%$ |
| CIFAR10 ResNet 8-15 | $94.39 \pm 5.52\%$ | $96.97 \pm 3.23\%$ | $98.96 \pm 1.43\%$ | $95.98 \pm 2.93\%$ |
| CIFAR10 ResNet 8 | $94.97 \pm 5.64\%$ | $99.13 \pm 1.94\%$ | $99.51 \pm 1.09\%$ | $96.01 \pm 2.84\%$ |

Table 1: Model comparison performing single-frame classification testing on the first frame in the time-lapse using downscaled versions of ResNet-8. The table shows species-specific accuracies and standard deviations across the predetermined train/test splits. The lowest species-specific accuracies are highlighted in light blue.

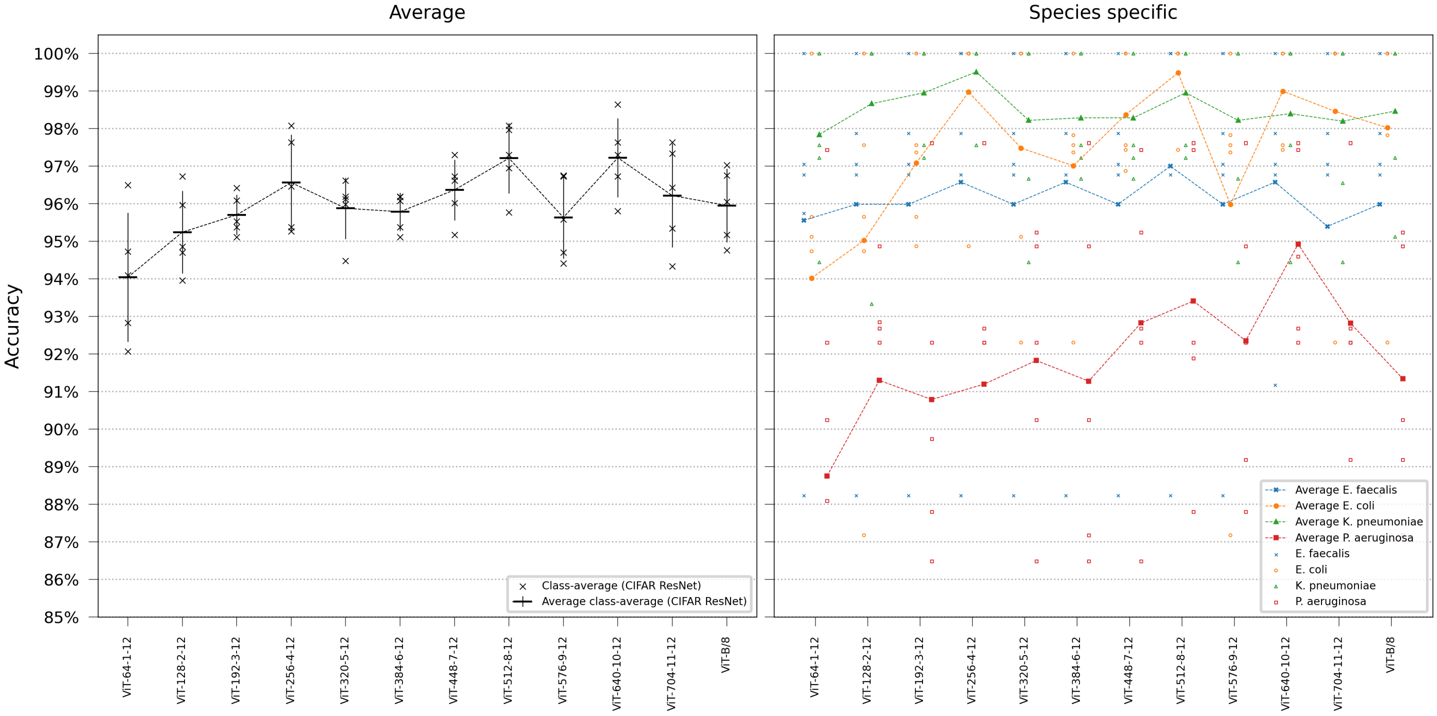


Figure 2: Model comparison performing single-frame classification of the first frame in the time-lapse using downscaled versions of ViT-B/8. Error bars represent the standard deviation in class-average accuracy from the five retrainings. Scatter plots depict the class-specific accuracy of all individual classifiers and the average class-specific accuracy over the five retrainings for each model. To reduce overplotting, a minor jitter was introduced along the categorical axis of the species-specific scatter plot. Lines are included not for interpolation or statistical inference purposes but to visually guide readers in tracking mean values on the ordinal scale.

| Model name | Enerococus facuialis | E coli | Klebsiella | Pseudamonas |
|---|---|---|---|---|
| ViT-64-1-12 | $95.56 \pm 4.39\%$ | $94.03 \pm 5.67\%$ | $97.85 \pm 2.31\%$ | $88.75 \pm 8.09\%$ |
| ViT-128-2-12 | $95.99 \pm 4.51\%$ | $95.03 \pm 4.83\%$ | $98.67 \pm 2.98\%$ | $91.30 \pm 4.32\%$ |
| ViT-192-3-12 | $95.99 \pm 4.51\%$ | $97.09 \pm 1.98\%$ | $98.96 \pm 1.43\%$ | $90.79 \pm 4.40\%$ |
| ViT-256-4-12 | $96.58 \pm 4.87\%$ | $98.97 \pm 2.29\%$ | $99.51 \pm 1.09\%$ | $91.20 \pm 6.09\%$ |
| ViT-320-5-12 | $95.99 \pm 4.51\%$ | $97.49 \pm 3.58\%$ | $98.22 \pm 2.56\%$ | $91.83 \pm 3.61\%$ |
| ViT-384-6-12 | $96.58 \pm 4.87\%$ | $97.01 \pm 2.84\%$ | $98.29 \pm 1.59\%$ | $91.28 \pm 4.85\%$ |
| ViT-448-7-12 | $95.99 \pm 4.51\%$ | $98.37 \pm 1.51\%$ | $98.29 \pm 1.59\%$ | $92.83 \pm 4.11\%$ |
| ViT-512-8-12 | $97.00 \pm 5.10\%$ | $99.49 \pm 1.15\%$ | $98.96 \pm 1.43\%$ | $93.41 \pm 4.15\%$ |
| ViT-576-9-12 | $95.99 \pm 4.51\%$ | $95.99 \pm 5.04\%$ | $98.22 \pm 2.56\%$ | $92.36 \pm 4.02\%$ |
| ViT-640-10-12 | $96.58 \pm 3.27\%$ | $99.00 \pm 1.37\%$ | $98.40 \pm 2.45\%$ | $94.93 \pm 2.53\%$ |
| ViT-704-11-12 | $95.40 \pm 5.79\%$ | $98.46 \pm 3.44\%$ | $98.20 \pm 2.58\%$ | $92.82 \pm 3.03\%$ |
| ViT-B/8 | $95.99 \pm 4.51\%$ | $98.03 \pm 3.33\%$ | $98.47 \pm 2.22\%$ | $91.34 \pm 3.56\%$ |

Table 2: Model comparison performing single-frame classification testing on the first frame in the time-lapse using downscaled versions of ViT/B-8. The table shows species-specific accuracies and standard deviations across the predetermined train/test splits. The lowest species-specific accuracies are highlighted in light blue.
